# SIEVEseq: One-stop differential expression, variability, and skewness analyses using RNA-Seq data

**DOI:** 10.1101/2024.04.09.588804

**Authors:** Hongxiang Li, Tsung Fei Khang

## Abstract

RNA-Seq data analysis is commonly biased towards detecting differentially expressed genes and insufficiently conveys the complexity of gene expression changes between biological conditions. This bias arises because discrete count models cannot fully and independently parameterize the mean, variance, and skewness of gene expression distributions. Therefore, a unified statistical framework that simultaneously tests differential expression, variability, and skewness is needed. We present SIEVEseq, a statistical methodology that provides such a framework. SIEVEseq embraces a compositional data analysis strategy to transform discrete RNA-Seq counts into continuous form with a distribution well-fitted by the skew-normal distribution. Both parametric and nonparametric simulations show that SIEVEseq better controls the false discovery rate and Type II error than existing differential expression methods. Analysis of the Mayo RNA-Seq dataset for Alzheimer’s disease demonstrates that gene sets with significant differences in mean, variance, and skewness between control and disease groups strongly predict disease state. Furthermore, functional enrichment analysis indicates that relying solely on differentially expressed genes identifies only part of the biological spectrum, whereas incorporating genes with differential variability and skewness reveals additional disease-related aspects. Cross-data and cross-methodology validation suggest the detected biological signals are genuine. The SIEVEseq R package and source codes are available at: https://github.com/Divo-Lee/SIEVEseq.

## 1 Introduction

The goal of gene expression analysis is to uncover genes with patterns of expression variation that are significantly associated with variation of some biological states. Three such patterns are of potential biological interest: (i) differential expression (DE); (ii) differential variability (DV); (iii) differential skewness (DS). These correspond to shifts in the mean, variance, and skewness of gene expression levels respectively when comparing two biological conditions.

The statistical detection of genes that are significantly up or down-regulated (i.e. DE genes) forms the main use of RNA-Seq data, but not all genes that are associated with change in biological states show change in mean expression level. For example, a gene’s previously tightly-controlled expression range can be lost, resulting in expression over a much wider range.^1,2^ Genes associated with modulation of the immune system, stress and hormonal regulation have high gene expression variability.^3^ Furthermore, gene expression variability often shows consistent differences between individuals with and without certain disease states.^4–6^ In cancer biology, cancer tissues show higher variability of gene expression compared with normal tissues.^6,7^ In all these cases, the focus is on genes that show differential variability (i.e. DV genes). Differential variability describes changes in the spread of expression values across samples and is often interpreted as a gain or loss of regulatory control, as well as increased or decreased inter-individual heterogeneity.^4^ Unfortunately, current methods of detecting DV genes are poorly integrated with the framework of commonly used DE tests. Differences in the skewness of gene expression levels have also been suggested to affect disease pathogenesis.^8–11^ Skewness characterizes the shape of the distribution of gene expression levels and potentially captures more nuanced patterns of variation compared to the mean and the variance. Differential skewness describes differences in the asymmetry of expression distributions in two groups, particularly those induced by presence of non-trivial extreme expression levels. Church et al. reported elevated positive skewness in the expression of genes within immune-related pathways when comparing cancer against control subjects.^10^ No methods are currently available for detecting the class of genes that show DS.

Conventional discrete count models have been effective in developing useful statistical tests of DE, but the lack of model parameters that explicitly characterize variance or skewness makes the development of DV and DS tests difficult. For context, we first provide an overview of DE and DV tests in RNA-Seq data analysis. The discrete negative binomial (NB) model underlies popular DE methods such as edgeR^12,13^ and DESeq2.^14^ For the NB model, the variance *σ*^2^ is modeled as a linear (*σ*^2^=ϕ*μ*) or quadratic (*σ*^2^=*μ*+ϕ*μ*^2^) function of the population mean *μ*, with ϕ as the dispersion parameter. These two parametrizations lead to the negative binomial 1 (NB1) and 2 (NB2) models, respectively. However, since mean and variance are confounded, it may be difficult to use these models effectively for analysis of gene expression variability.^15^ Other models such as the Generalized Poisson^16^ and the Poisson-Tweedie distribution^17^ have also been used. Alternatively, the voom algorithm^18^ assumes a Gaussian distribution for the logarithm of RNA-Seq counts, and repurposes the limma pipeline^19^ used in microarrays for DE analysis. Gauthier et al. proposed dearseq,^20^ a distribution-agnostic method that uses a test statistic based on the variance component score. Nonparametric methods (NOISeq^21,22^ and SAMSeq^23^) are available, but they are less commonly used compared to DESeq2 or edgeR. Nevertheless, for large sample sizes, the Wilcoxon rank-sum test seems superior to edgeR, DESeq2 and dearseq with respect to the false discovery rate (FDR).^24^

In contrast, only five DV methods are available: (i) DiffVar,^25^ an empirical Bayes method that uses the limma framework;^19^ (ii) MDSeq,^26^ which uses an NB1 generalized linear model to test DE and DV separately; (iii) the generalized additive model for location, scale and shape (GAMLSS),^27^ which tests the effects of biological factors on a gene’s Poisson and non-Poisson variation using the NB2 model;^28^ (iv) DiffDist,^29^ a hierarchical Bayesian model based on the NB2 distribution; (v) clrDV,^30^ our contribution that uses the compositional data framework in the present work but restricted to DV test.

The non-normality of gene expression distribution, particularly as characterized by skewness, may be potentially crucial under certain biological contexts.^11^ It can shed light on gene functionalities that cannot be discovered using conventional analysis of difference in mean - and, less commonly, differences in variability.^4,5,31^ The relevance of skewness of distributions of gene expression as measured using microarray data was documented in several studies.^8,9,32,33^ More recently, Church et al. highlighted skewness as an effective indicator of biological heterogeneity in gene pathways associated with specific cancer cohorts.^10^ Despite these studies, no methods are currently available for testing DS. A unified framework that enables the simultaneous testing of mean, variance and skewness requires a model where these parameters are explicit and independent of one another. This suggests transformation of RNA-Seq count data into continuous data, followed by their appropriate modeling using a continuous distribution. Quinn et al. adopted a compositional data framework for RNA-Seq data and repurposed ALDEx2^34,35^ - originally developed for differential abundance testing in microbiome data - for detecting DE genes.^36^ ALDEx2 incorporates a Dirichlet process to produce centered log-ratio (CLR) transformation of count data.^37^ The transformed data is assumed Gaussian, and a Welch t-test or Wilcoxon test is applied to test for differential abundance between two populations. Interestingly, ALDEx2 has been shown to be competitive against edgeR and DESeq2.^36^ Although the Welch t-test is robust against non-Gaussian data,^38^ without explicitly modeling the CLR-transformed data, it is unclear how tests concerning aspects such as equality of variances or skewness can be done.

Toward the goal of a unified framework for RNA-Seq data analysis, we need to use a model that is both mathematically tractable and provides a good empirical fit the transformed RNA-Seq data. Our solution is SIEVEseq (one-*S*top d*I*fferential *E*xpression, *V*ariability, and sk*E*wness analyses using RNA-*Seq* data), which can perform simultaneous testing of differential expression, variability, and skewness of genes using RNA-Seq data. SIEVEseq adopts a compositional data analysis approach to modeling discrete RNA-Seq count data, applies Aitchison’s CLR transformation^37^ to convert them into continuous form, and uses the skew-normal distribution^39^ to model them. With SIEVEseq, genes with significant variation in skewness can now be detected for the first time, alongside those with significant variation in mean and variance.

## 2. Materials and Methods

### 2.1 The compositional data framework and the centered log-ratio transformation

Let *X _gi_* be the read count of gene *g* and sample *i*, where *g* = 1,2, …, and *i* = 1,2, …,. The *i*th sample data becomes compositional if we normalize the counts by dividing them with the sum of gene counts. Application of the CLR transformation converts the variable space from the simplex to the Euclidean space, for which standard mathematical and statistical methods operate in. The CLR-transformation of the count of the *g*th gene is defined as the logarithm of the gene count divided by the geometric mean of gene counts across all *G* genes. To avoid multiplication or division by 0, a pseudo-value of 0.5 replaces any zero gene counts.

### 2.2 Modeling Centered Log-ratio-transformed Data Using the Skew-normal Distribution

Denote the CLR-transformed count from gene *g* in sample *i* by *Y_gi_*. We model the random variable *Y_gi_* using a skew-normal distribution with centered parameters (CP). Notationally, we write *Y_gi_* ∼ SN_C_(*μ_g_*,*σ_g_*,*γ_g_*), where *μ_g_* is the mean, *σ_g_* is the standard deviation, and *γ_g_* is the skewness parameter (see Supplementary Materials S1 for mathematical details). The parameter vector 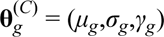 has parameter space R×R^+^×(-*k*,*k*), where 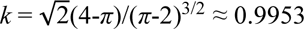. The skew- normal distribution simplifies to the normal distribution with mean *μ_g_* and variance 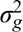 when *γ* = 0. Given two populations and sufficiently large sample sizes, tests of equality of mean, standard deviation and skewness can be done using the Wald statistic:

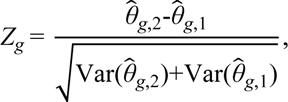

where *g* = 1,2,…,*G* index the genes, and *j* = 1,2 index the groups. Denote the maximum likelihood estimator (MLE) or the maximum penalized likelihood estimator (MPLE)^40,41^ as 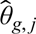. Then, for testing DE, 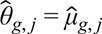; for testing DV, 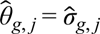; for testing DS, 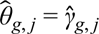. For estimating 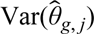, the inverse of the Fisher information matrix of 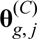 is used. When sample size is sufficiently large, *Z_g_* converges to the standard normal distribution. The Benjamini-Yekutieli (BY) procedure,^42^ which allows for arbitrary dependence between the tested hypotheses, is used to control the FDR and adjust *p*-values for multiple hypothesis testing.

Under the SIEVEseq framework, eight gene classes are possible (see also Supplementary Fig. S1). These range from genes that do not show DE, DV or DS, to genes that simultaneously show them (Table 1).

**Table 1.** Eight gene classes identifiable using SIEVEseq. Abbreviations: DE = differential expression; DV = differential variability; DS = differential skewness; 1 = True; 0 = False.

| Gene class | DE | DV | DS | Label |
| --- | --- | --- | --- | --- |
| non-DE, non-DV and non-DS | 0 | 0 | 0 | 000 |
| Pure DE | 1 | 0 | 0 | 100 |
| Pure DV | 0 | 1 | 0 | 010 |
| Pure DS | 0 | 0 | 1 | 001 |
| DE, DV, non-DS (DEV) | 1 | 1 | 0 | 110 |
| DE, non-DV, DS (DES) | 1 | 0 | 1 | 101 |
| non-DE, DV, DS (DVS) | 0 | 1 | 1 | 011 |
| DE, DV, DS (DEVS) | 1 | 1 | 1 | 111 |

### 2.3 Data Description and Preprocessing

To assess the suitability of the skew-normal distribution for fitting CLR-transformed counts and to compare SIEVEseq’s performance with existing methods for DE tests, with a particular focus on controlling the FDR and the probability of Type II error, it is essential to simulate RNA-Seq count data distributions using both realistic parameter values and a nonparametric resampling strategy. For this purpose, we used three real RNA-Seq datasets with varying sample sizes to facilitate reliable parameter estimation of the NB2 model. Two datasets with sufficiently large sample sizes in each group were used to enable a nonparametric resampling–based simulation design.

The first dataset (GEO accession number: GSE123658) contains whole blood RNA-Seq data from 39 Type 1 diabetes patients and 43 healthy donors, with 16,785 genes.^43^ The second dataset (GEO accession number: GSE150318) consists of longitudinal RNA-Seq data from 114 short-lived killfish *Nothobranchius furzeri*,^44^ with samples collected at 10 and 20 weeks of age, covering 26,739 genes. The third dataset (GEO accession number: GSE179250) comprises 29,245 genes from 192 human liver samples.^45^ The fourth dataset (GEO accession number: GSE229705) comprises RNA-Seq profiles of 60,591 genes across 123 paired tumor and normal samples.^46^ The fifth dataset (GEO accession number: GSE150910) consists of RNA-Seq profiles of 18,838 genes measured in whole-lung tissues from 82 chronic hypersensitivity pneumonitis (CHP) patients, 103 idiopathic pulmonary fibrosis (IPF) patients, and 103 unaffected controls.^47^ These datasets will subsequently be referred to as the Valentim, Kelmer, Zhou, Dolgalev, and Furusawa datasets, respectively.

For empirical assessment, we used the Mayo RNA-Seq dataset,^48^ which contains 278 samples and 64,253 transcripts. Access to this dataset was obtained from the AD Knowledge Portal (https://adknowledgeportal.synapse.org) after obtaining permission from the database owner (1 September 2022; ID: 9603055). This dataset comprises RNA isolated from the temporal cortex of patients with four biological conditions: control (*n*=80), Alzheimer’s disease (AD; *n*=84), progressive supranuclear palsy (*n*=80), and pathologic aging (*n*=30). Here, we compared the AD group against the control group. We chose the Mayo RNA-Seq dataset for three main reasons. Firstly, the study has a rigorous experimental design and quality control for ensuring data fidelity. Secondly, the sample size per group is sufficiently large (50 or more per group) to allow the standard deviation and skewness parameters of the skew-normal model to be reliably estimated.^40,49^ Thirdly, analysis of an AD dataset enables us to leverage on substantial literature to cross-check the biological significance of genes detected using SIEVEseq.

For cross-data validation of gene ontology (GO) biological processes identified from the Mayo RNA-Seq dataset, we used the Nakayama dataset^50^ (GEO accession number: GSE249477) as an external cohort. This dataset comprises whole blood RNA-Seq profiles of 21,479 genes from a cohort of 62 subjects from Japan. The study participants were clinically categorized into three groups: healthy controls (*n*=21), patients with mild cognitive impairment (MCI; *n*=20) due to AD, and patients with AD (*n*=21). We focused on the 21 AD patients and 21 controls to provide a direct comparison with the primary Mayo cohort.

To filter genes, those with an average CPM ≤ 0.5 or with zero counts in at least 85% of the samples were excluded. After this step, the Valentim, Kelmer, Zhou, Dolgalev, and Furusawa datasets retained 12,283, 16,670, 17,862, 20,151, and 14,316 genes, respectively. In the Mayo RNA-Seq dataset, after removing samples with missing class labels, 78 and 82 samples remained for the control and AD groups, respectively. Similarly, in the Nakayama dataset, 13,078 genes remained for the AD-control comparison after handling missing values and gene filtering.

For SIEVEseq and ALDEx2, the CLR transformation was applied to the raw counts. For the eight DE methods (edgeR, DESeq2, voom, tweeDEseq, NOISeq, DSS, dearseq, and Wilcoxon rank-sum test), raw counts were normalized using the trimmed mean of M values (TMM) approach.^51^

### 2.4 Parametric Simulation Design for Evaluating Performance of Differential Expression Tests

To compare the performance of SIEVEseq with existing DE methods, RNA-Seq count data were simulated from an NB2 model with model parameters estimated from real datasets using polyester^52^ R. Three RNA-Seq datasets with reasonably large sample sizes were used to facilitate reliable estimation of the standard deviation and skewness parameters.

Assuming the distribution of the RNA-Seq count data follows the NB2 model, we randomly sampled 5000 of the filtered genes and then estimated the parameters of the NB2 model using samples from one group (Type 1 diabetes group in the Valentim dataset (*n* = 39); 10-weeks group in the Kelmer dataset (*n* = 114); all liver samples in the Zhou dataset (*n* = 192)). After this, we generated 200 biological replicates from the estimated NB2 models. The goodness-of-fit of the skew-normal model on the distribution of the CLR-transformed simulated count data was checked using the Kolmogorov-Smirnov (KS) test.

Using the FDR and the probability of Type II error (*β*) as performance metrics (see Supplementary Materials S1.5), we compared SIEVEseq against nine other DE methods: edgeR, DESeq2, voom, DSS, tweeDEseq, NOISeq, ALDEx2, dearseq, and the Wilcoxon rank-sum test. We chose edgeR, DESeq2 and voom because of their popularity as DE tests. Then, to comprehensively survey the landscape of DE tests using a diversity of approaches, we included two nonparametric methods: NOISeq and the Wilcoxon rank-sum test, as well as DSS, dearseq and tweeDEseq, which are based on ideas of dispersion shrinkage, variance component score, and the Poisson-Tweedie model, respectively. RNA-Seq data were similarly simulated using the same three datasets with polyester^50^ R. For each of the three datasets, a total of 2,000 filtered genes were randomly selected, and their mean (*μ*) and size (1/ϕ) parameters under assumption of an NB2 model were estimated. The samples used were the 39 Type I diabetes patients for the Valentim dataset, the 114 fish at 10 weeks of age for the Kelmer dataset, and all 192 livers for the Zhou dataset.

In a two-group comparison setting, both groups have identical NB2 model parameters under the null hypothesis of equal mean. Hence, to simulate DE genes, we spiked 10% of the genes in one of the two groups to be differentially expressed, approximately half of which are up-regulated and the remainder down-regulated. This was done by multiplying or dividing their estimated mean parameter with a random value uniformly chosen between 2 to 4. Biological replicates of size 30, 50, and 100 per group were generated from the NB2 model, with 30 instances simulated for all three datasets.

We defined the region of desirable performance as where both FDR and *β* are below 0.05. Then, we computed the percentage of simulated instances falling within this region (*θ*). We identified genes with adjusted *p*-value below 0.05 as DE genes, and recorded the computing time of each method.

To evaluate performance similarity of DE methods, we used a cluster heatmap with the Ward clustering algorithm and Euclidean distance for between-method distances. We computed the mean and standard deviation of FDR and *β* (used as features) for the 30 simulated instances across each of the three sample size scenarios, and subjected them to min-max normalization before constructing the cluster heatmap.

Finally, to evaluate the effect of using pseudo-values that are smaller than 0.5, we conducted a sensitivity analysis using two smaller pseudo-values (0.01 and 0.1).

### 2.5 Nonparametric Simulation Design for Evaluating Performance of Differential Expression Tests

To further validate whether the superior performance of SIEVEseq persists when the underlying distribution deviates from the NB assumption, we implemented a nonparametric simulation study using the SimSeq^53^ R package. This approach constructs simulated counts by resampling from the empirical distributions of a real dataset, and captures the empirical variance and skewness inherent in biological samples. To obtain moderately large sample sizes in the simulated conditions, we used two RNA-Seq datasets (Dolgalev dataset - 123 paired tumor-normal samples; Furusawa dataset - 103 IPFs vs. 103 controls) with sufficiently large sample sizes in each group.

We randomly selected 2,000 filtered genes for each simulation pool. Following the SimSeq protocol, 10% of these genes were spiked to become DE genes with an absolute log fold-change greater than 1. Under this nonparametric framework, biological replicates were generated for two sample sizes (30 and 50 per group), with 30 independent instances simulated for each scenario. We restricted the maximum sample size to 50 to satisfy the SimSeq operation contraint,2*n*≤max{*N*_1_,*N*_2_}, where *N*_1_ and *N*_2_ are the sample sizes in the two respective groups of the source dateset.

Consistent with the design of our parametric simulation evaluation, we also used FDR and *β* to assess model performance. We compared SIEVEseq against eight other DE methods: edgeR, DESeq2, voom, DSS, tweeDEseq, NOISeq, ALDEx2, and dearseq. The Wilcoxon rank-sum test was excluded to avoid circular bias, as the SimSeq algorithm itself applies this test to define the ground-truth DE genes. We defined the region (*θ*) of desirable performance where FDR < 0.05 and *β*<0.2, and subsequently computed the percentage of simulated instances falling within this region. Genes identified as DE genes with an adjusted p-value below 0.05. Finally, the similarity of DE methods was evaluated using the same cluster heatmap analysis and hierarchical clustering parameters as described in the parametric simulation study.

### 2.6 Empirical Validation

We applied SIEVEseq to the Mayo RNA-Seq dataset to assess its capability for detecting DE, DV, and DS genes that are contextually meaningful in Alzheimer’s disease (AD). The KS test was used to check the fit of the skew-normal distribution on the CLR-transformed count data.

For comparison of DE methods, we selected those that showed reasonably good performance in the simulation study. Comparison of DV methods were reported previously by us.^30^ Volcano plots were used to inspect joint patterns of biological and statistical significance. Biological significance was measured using the difference of mean for DE genes, log_2_ of the ratio of standard deviation for DV genes, and the difference of skewness for DS genes. Thus, zero corresponds to genes with no significant biological difference between the two groups. Significant DE, DV and DS genes were flagged using the significance score method (see Supplementary Materials S2).^54^ Sets of DE genes detected using the different methods were examined using Venn diagrams. Finally, the joint distributions of estimated mean, standard deviation, and skewness parameters for the control and the AD group were visualized using contour heatmaps.

Subsets of the genes detected using SIEVEseq that are strongly predictive of the AD state were identified using the Generalized, Unbiased Interaction Detection and Estimation (GUIDE Version 40.3) decision tree algorithm.^55,56^ To assess the informativeness of the set of significant DE, DV, and DS genes (the top 10 genes on the left and the right tails of the distribution of biological significance scores) for predicting the AD state, we estimated the out-of-bag generalization error using the GUIDE random forests model with default hyperparameters (equal priors, unit misclassification costs, univariate split highest priority, no interaction splits, number of trees = 1000, SE parameter = 0.25). For benchmarking, AD state classification using GUIDE was done using a random subset of 60 putatively uninformative genes that are non-DE, non-DV, and non-DS with BY-adjusted *p*-value of 1.

Gene set functional enrichment analysis was done using WebGestalt.^57,58^ We explored enriched biological processes in the GO functional database using over representation analysis (ORA) of the following five gene lists: (i) high confidence DE genes obtained the intersection of sets of DE genes detected using SIEVEseq, edgeR, DESeq2, voom, tweeDEseq, ALDEx2, DSS and Wilcoxon; (ii) non-DE, DV and/or DS genes; (iii) pure DV genes; (iv) pure DS genes; and (v) the union of DE, DV, and DS genes. The human genome reference set was chosen. Only the top 20 GO terms were included, and affinity propagation was used for redundancy reduction. GO hierarchies were checked using WebGestalt’s GOView application.^57^

To improve confidence in the reliability of biological processes inferred using the DE, DV and DS genes in the Mayo RNA-Seq dataset, we performed additional cross-methodological and cross-data validations. We applied Gene Set Enrichment Analysis^59^ (GSEA) on the Nakayama dataset to detect GO biological processes that are significantly enriched. Recovering a substantial proportion of significant GO biological processes across independent AD cohorts using GSEA provides cross-methodological support that the DE, DV and DS genes detected using SIEVEseq carry meaningful biological signals. To this end, we constructed a composite ranking statistic *U* that accentuates the most prominent distributional divergence of each gene in Nakayama dataset. Specifically, we assigned each gene a composite score based on the difference of means 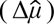, the log2 fold-change of the standard deviation ratio 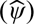, and the difference of skewness (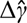; see 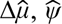, and 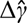 in Supplementary Materials Section S2). Thus, we defined 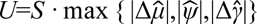, where *S* is the sign of the component with the largest magnitude. This statistic prioritizes genes exhibiting the largest shift in any of the three distributional moments, while preserving the direction of the effect.

### 2.7 Computing Tools and Environment

Computational work for DE, DV and DS tests was done using a MacBook Pro (macOS Monterey version 12.5) with 32GB RAM, an M1 pro chip and a 10-core CPU. The R (version 4.2.1)^60^ computing environment was used. Supplementary Materials S6 gives the complete list of R packages used. For converting ENSEMBL gene ID to gene symbol, we used the application programming interface of BioTools.fr.^61^

## 3. Results

### 3.1 Parametric Simulation Study

The skew-normal distribution fits the CLR-transformed count data of genes in the three datasets well, with 99.2%, 98.9% and 98.7% of genes having KS-test *p* -values above 0.05 (Fig. 1). Graphical summaries of the parametric simulation results are given in Fig. 2, 3, and 4. For brevity, details of the summary statistics of the performance metrics are given only for the Valentim dataset (Table 2). For the other two simulations, see Supplementary Tables S1 and S2.

**Fig. 1.**
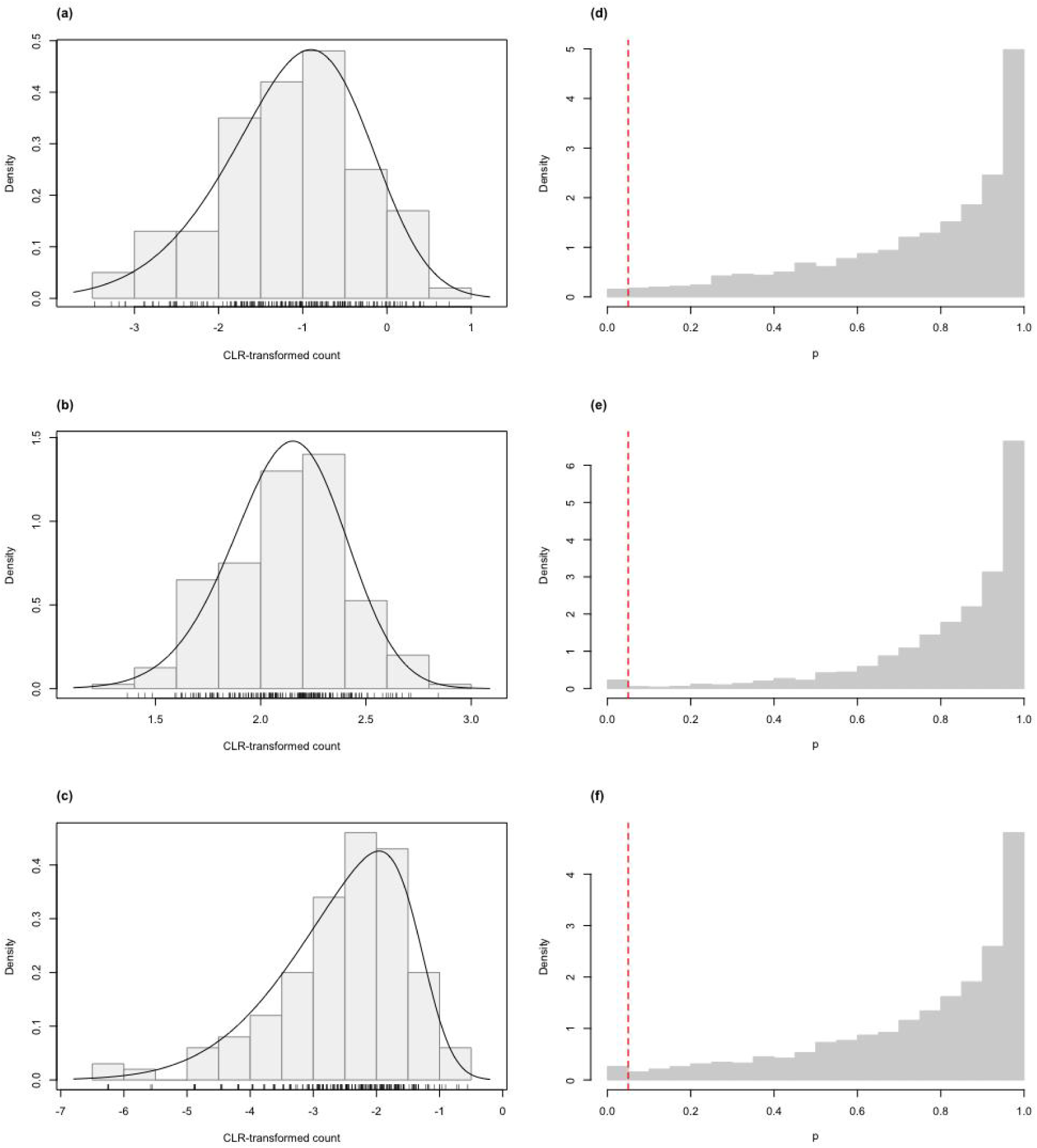
Histograms of CLR-transformed counts for three selected genes with fitted skew-normal curve for (a) the Valentim dataset (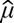=-1.256 (s.e.=0.061), 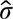=0.861 (s.e.=0.046) and 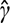=-0.450 (s.e.=0.172)); (b) the Kelmer dataset (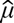=2.119 (s.e.=0.019), 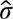=0.274 (s.e.=0.014) and 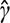=-0.232 (s.e.=0.196)); (c) the Zhou dataset (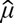=-2.564 (s.e.=0.073), 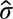=1.040 (s.e.=0.057) and 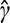=-0.799 (s.e.=0.071)). Distribution of the p-values of the Kolmogorov-Smirnov goodness-of-fit tests of the skew-normal model for genes in the simulated (d) Valentim dataset; (e) Kelmer dataset; and (f) Zhou dataset. The skew-normal model gives good fit to about 99.2%, 98.9%, and 98.7% of the genes in (d), (e), and (f), respectively. The red dashed line corresponds to the threshold p-value of 0.05.

**Fig. 2.**
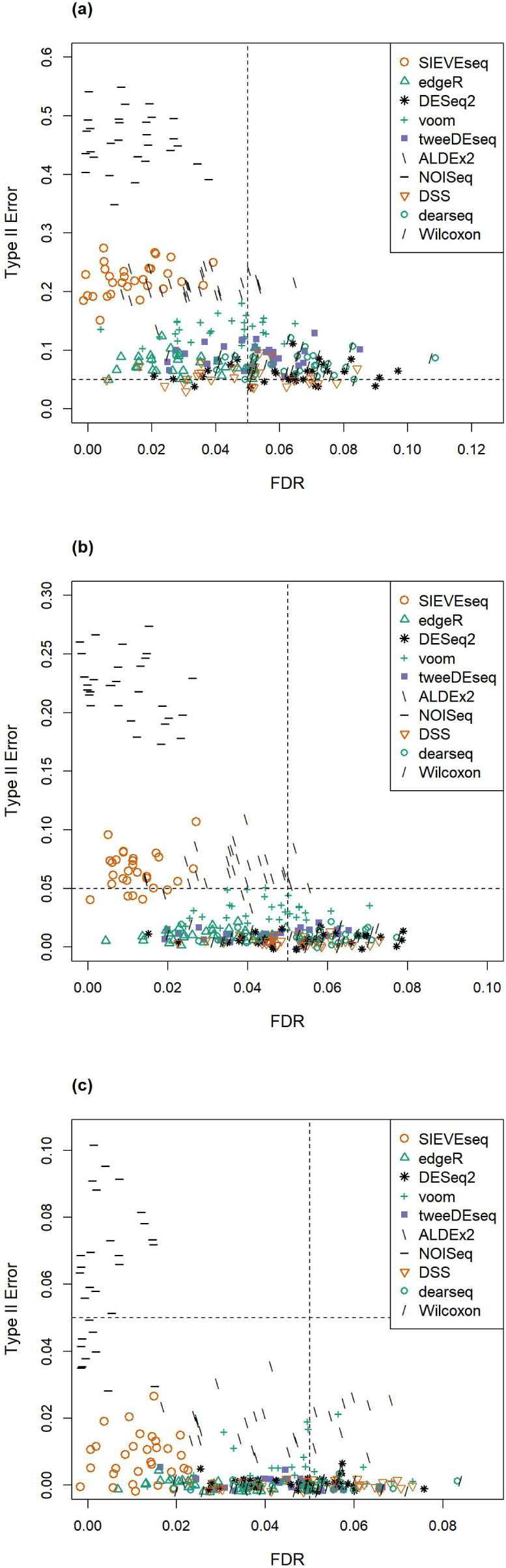
Scatter plots of the probability of Type II error (*β*) against FDR for simulated data from the Valentim dataset (30 instances) for three sample size per group scenarios: (a) 30; (b) 50; (c) 100. Dashed lines represent the desired threshold FDR and *β* of 0.05.

**Fig. 3.**
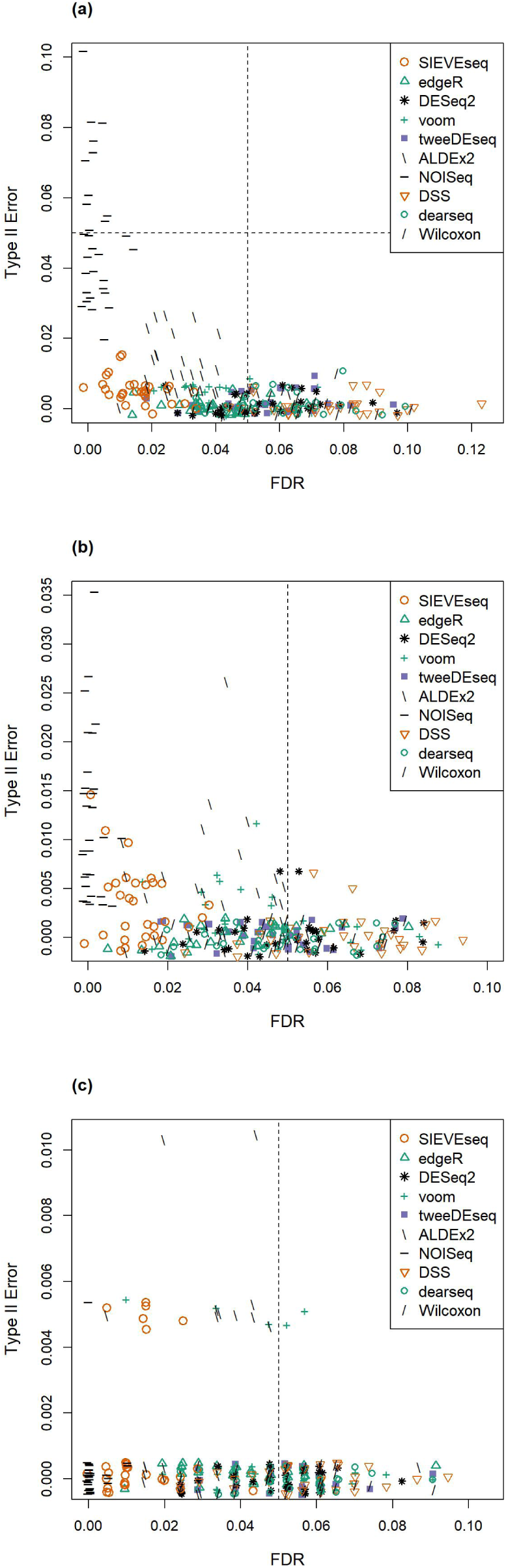
Scatter plots of the probability of Type II error (*β*) against FDR for simulated data from the Kelmer dataset (30 instances) for three sample size per group scenarios: (a) 30; (b) 50; (c) 100. Dashed lines represent the desired threshold FDR and *β* of 0.05.

**Fig. 4.**
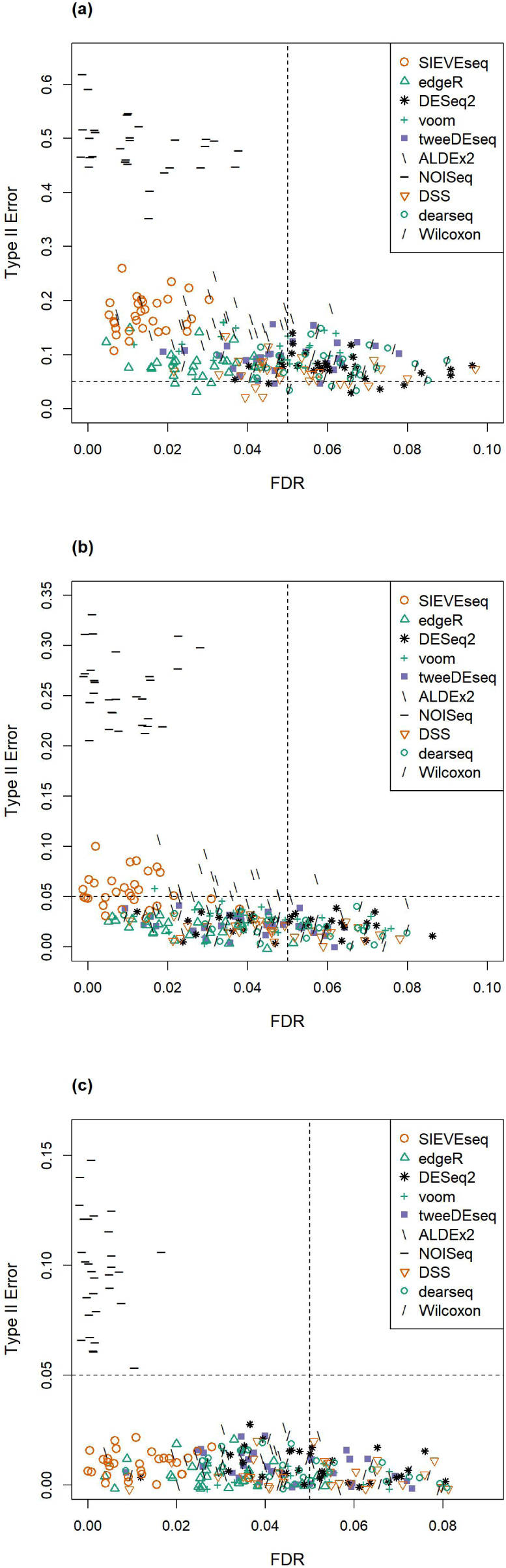
Scatter plots of the probability of Type II error (*β*) against FDR for simulated data from the Zhou dataset (30 instances) for three sample size per group scenarios: (a) 30; (b) 50; (c) 100. Dashed lines represent the desired threshold FDR and *β* of 0.05.

**Table 2.** The mean of false discovery rate (FDR*), mean probability of Type II error (*β*^†^), and percentage of simulated instances with FDR < 0.05 and *β* <0.05 (*θ*^‡^) for each of the ten DE tests applied to data simulated from the Valentim dataset (30 instances) at three different sample sizes. Standard deviation in parentheses.

| Method | Sample size per group |  |  |
| --- | --- | --- | --- |
|  | 30 | 50 | 100 |
| SIEVEseq | 0.014 (0.011) | 0.012 (0.006) | 0.012 (0.007)* |
|  | 0.221 (0.028) | 0.066 (0.015) | 0.008 (0.007)† |
|  | 0% | 13.3% | 100%‡ |
| ALDEX2 | 0.034 (0.014) | 0.038 (0.011) | 0.041 (0.015) |
|  | 0.210 (0.022) | 0.066 (0.016) | 0.017 (0.008) |
|  | 0% | 10% | 66.7% |
| NOISeq | 0.013 (0.011) | 0.009 (0.008) | 0.004 (0.005) |
|  | 0.458 (0.047) | 0.223 (0.026) | 0.062 (0.020) |
|  | 0% | 0% | 30% |
| edgeR | 0.029 (0.013) | 0.031 (0.012) | 0.029 (0.010) |
|  | 0.076 (0.018) | 0.010 (0.005) | 0.000 (0.001) |
|  | 0% | 96.7% | 100% |
| DESeq2 | 0.062 (0.019) | 0.054 (0.016) | 0.047 (0.012) |
|  | 0.060 (0.017) | 0.007 (0.005) | 0.000 (0.002) |
|  | 6.7% | 40% | 50% |
| voom | 0.062 (0.019) | 0.045 (0.012) | 0.046 (0.014) |
|  | 0.134 (0.021) | 0.031 (0.010) | 0.004 (0.006) |
|  | 0% | 60% | 66.7% |
| DSS | 0.049 (0.018) | 0.051 (0.011) | 0.051 (0.015) |
|  | 0.056 (0.015) | 0.004 (0.004) | 0.000 (0.000) |
|  | 13.3% | 46.7% | 46.7% |
| dearseq | 0.063 (0.016) | 0.053 (0.015) | 0.047 (0.014) |
|  | 0.077 (0.017) | 0.009 (0.005) | 0.000 (0.001) |
|  | 0% | 33.3% | 50% |
| Wilcoxon | 0.047 (0.012) | 0.045 (0.012) | 0.045 (0.014) |
|  | 0.123 (0.024) | 0.022 (0.009) | 0.001 (0.003) |
|  | 0% | 76.7% | 73.3% |
| tweeDEseq | 0.052 (0.014) | 0.046 (0.014) | 0.044 (0.012) |
|  | 0.089 (0.019) | 0.010 (0.005) | 0.000 (0.001) |
|  | 0% | 56.7% | 73.3% |

Overall, SIEVEseq, ALDEx2 and NOISeq prioritise FDR control over *β*. In all three simulations, these three methods, along with edgeR, are the top four methods with lower mean FDR. Specifically, NOISeq generally has the lowest mean FDR, followed by SIEVEseq, edgeR, and ALDEx2 in the simulated Valentim and Zhou datasets. In the simulated Kelmer dataset, NOISeq again has the lowest FDR, followed by SIEVEseq, ALDEx2, and edgeR. However, we note that NOISeq also has the highest mean *β* among all DE methods compared. ALDEx2 has relatively higher mean *β*, followed by SIEVEseq and edgeR. In all three simulations, edgeR has mean FDR that is relatively constant at about 0.03 (range: 0.026 to 0.042) across the three sample size scenarios. Furthermore, its mean *β* is at most about 0.08 at *n*=30, and almost 0 as *n* increases to 100. In contrast, DESeq2, voom, tweeDEseq, DSS, dearseq, and the Wilcoxon rank-sum test control FDR at around 0.05, with generally greater variation than edgeR’s. All methods have decreasing *β* as sample size increases.

At *n*=100, only SIEVEseq has *θ*=100% for all three simulations (see Tables 2, S1, and S2). SIEVEseq, edgeR, and ALDEx2 have *θ* ranging from 66.7-100%, 70-100% and 83.3-100% for the three simulations, compared to 46.7-73.3%, 26.7-67.7%, and 43.3-66.7% for DESeq2, voom, tweeDEseq, DSS, dearseq, and the Wilcoxon rank-sum test. NOISeq shows inconsistent performance, with *θ* ranging from 0% to 100%.

The cluster heatmap (Fig. 5) shows that NOISeq has mean *β* that is consistently the highest, in contrast with its mean FDR, which is consistently the lowest. SIEVEseq is similar to NOISeq with respect to mean FDR performance, but excels with substantially lower mean *β*. This suggests that SIEVEseq maintains the conservativeness of NOISeq without excessive compromise in *β*. Conversely, while other methods often have lower mean *β* values, their mean FDR values are correspondingly higher.

**Fig. 5.**
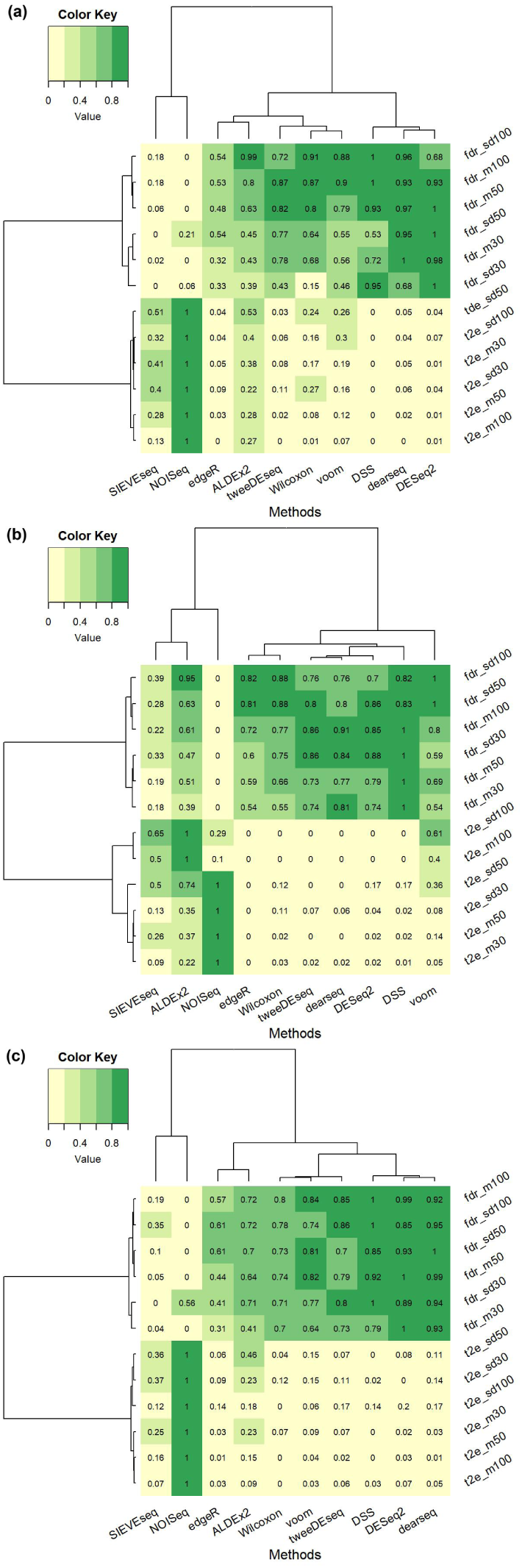
Cluster heatmaps for showing similarity of DE methods (columns) with respect to summary statistics (rows) of FDR and probability of Type II error (*β*) at three sample size scenarios (*n*=30,50,100) for the (a) Valentim dataset; (b) Kelmer dataset; and (c) Zhou dataset. The number in a cell represents the mean (over 30 instances) of the summary statistic of a specific feature. Abbreviations: fdr = FDR; m = mean; sd = standard deviation; t2e = probability of Type II error; numerical suffixes indicate corresponding sample size scenario.

SIEVEseq took longer time to run for small sample sizes, but its speed improved for larger sample sizes (Supplementary Table S3). At *n*=100 for all three simulations, edgeR, DESeq2, voom, DSS, dearseq, and the Wilcoxon rank-sum took about 5 s or less to complete, whereas NOISeq, ALDEx2 and SIEVEseq took about 11 to 46s. The slowest method was tweeDEseq, which was 14 to 17 times slower than the second (ALDEx2) and the third (SIEVEseq) slowest methods, respectively.

The result of the sensitivity analysis for different choice of pseudo-values shows that 0.5 generally provides better control of both FDR and *β*, compared to the smaller pseudo-values of 0.01 and 0.1, for data simulated from the Valentim, Kelmer, and Zhou datasets across three sample size scenarios (Supplementary Table S4; Supplementary Figs. S2-S4). Specificaly, at sample size of 30, all three pseudo-values give very similar *β* values for all the simulated datasets. For moderately large sample size of 50, using 0.5 as the pseudo-value tends to provide uniformly better control of both FDR and *β*.

### 3.2 Nonparametric Simulation Study

Graphical summaries of the nonparameteric simulation results are given in Fig. 6. For brevity, details of the summary statistics of the performance metrics for the Dolgalev and Furusawa datasets are given in Supplementary Tables S5 and S6, respectively. In these scenarios, SIEVEseq, ALDEx2, and NOISeq demonstrated superior FDR control compared to conventional parametric methods. SIEVEseq maintained a low mean FDR across all conditions (range: 0.012 to 0.025). In contrast, widely used methods such as edgeR, DESeq2, and DSS failed to control the FDR within the nominal 0.05 level, with mean FDRs reaching as high as 0.231 for edgeR for the Dolgalev dataset simulation (= 50). NOISeq maintained the most stringent FDR control in some cases, but it consistently exhibited the highest β among all compared methods (e.g. β= 0.749 at = 30 for the Dolgalev dataset), indicating low power in detecting true DE genes. SIEVEseq and ALDEx2 provided a better balance, with SIEVEseq consistently achieving lowerβ values than ALDEx2 across all scenarios.

**Fig. 6.**
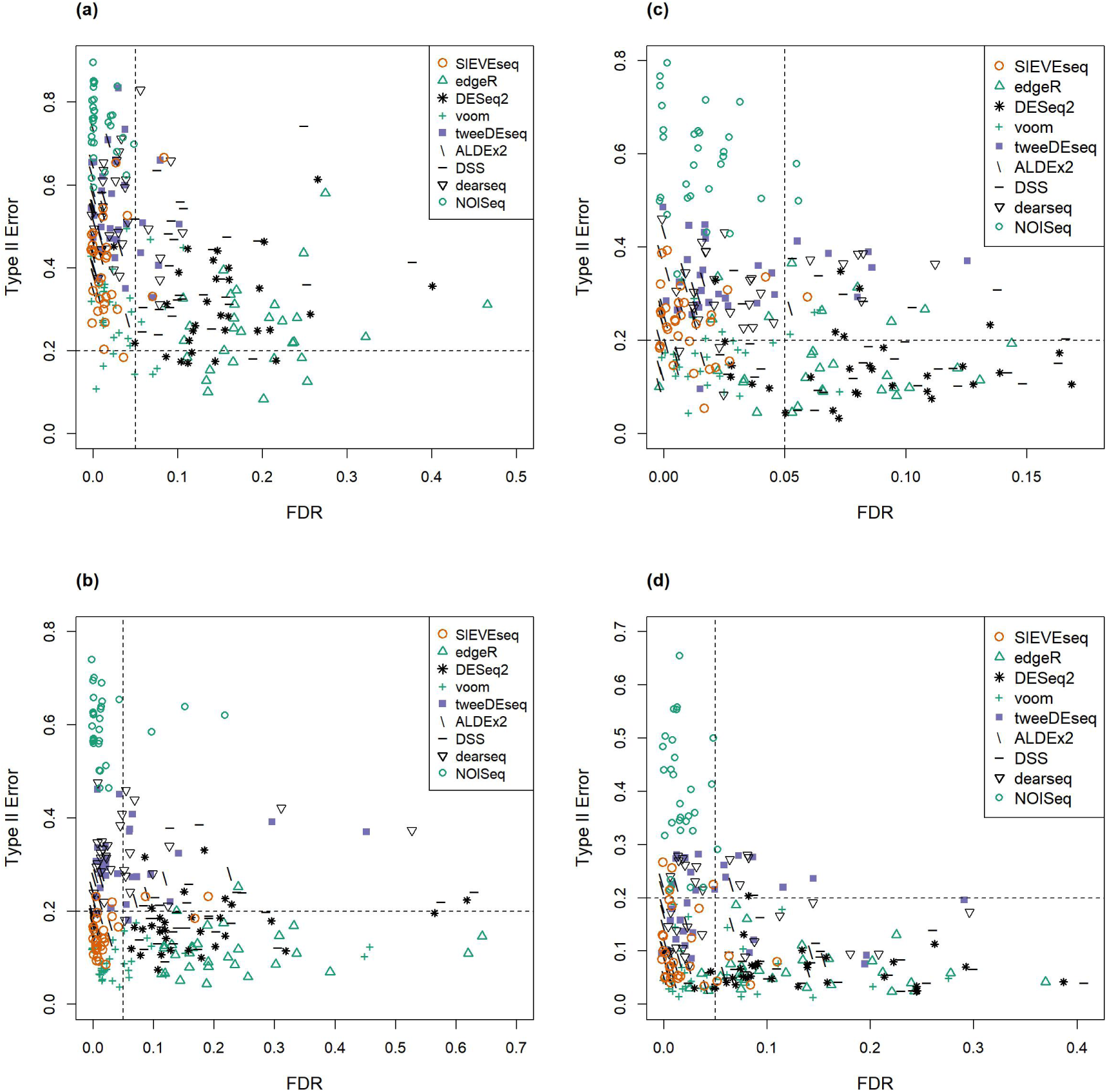
Scatter plots of the probability of Type II error (*β*) against FDR for simulated data from the: (a) Dolgalev dataset (30 instances) with a sample size of 30 per group; (b) Dolgalev dataset (30 instances) with a sample size of 50 per group; (c) Furusawa dataset (30 instances) with a sample size of 30 per group; (d) Furusawa dataset (30 instances) with a sample size of 50 per group. The vertical and horizontal dashed lines represent the target FDR threshold of 0.05 and the target *β* threshold of 0.2, respectively.

SIEVEseq was the most reliable method with θ= 83.3%, whereas for most other methods, including edgeR and DESeq2, dropped to 0%. Similarly, in the Furusawa dataset simulation with = 50, SIEVEseq achieved the highest θ of 76.7%. These results suggest that SIEVEseq is comparatively more robust against complexities of RNA-Seq data distributions with moderately large sample sizes byproviding stable FDR control and maintaining competitive β under non-canonical models of count data distribution.

The cluster heatmaps (Fig. 7) show that edgeR, DESeq2, and DSS cluster closely together. These three methods had the highest mean and standard deviation of FDR, which indicate a failure to control false discoveries. In contrast, SIEVEseq and ALDEx2 had consistently similar low mean FDR levels. NOISeq achieved the lowest mean FDR, but its β remained the highest across both datasets. Interestingly, SIEVEseq matched the low FDR performance of NOISeq but excelled with substantially lower mean β. Thus, it seems that SIEVEseq successfully maintains the conservativeness of NOISeq without an excessive increase inβ, even in a nonparametric simulation scenario.

**Fig. 7.**
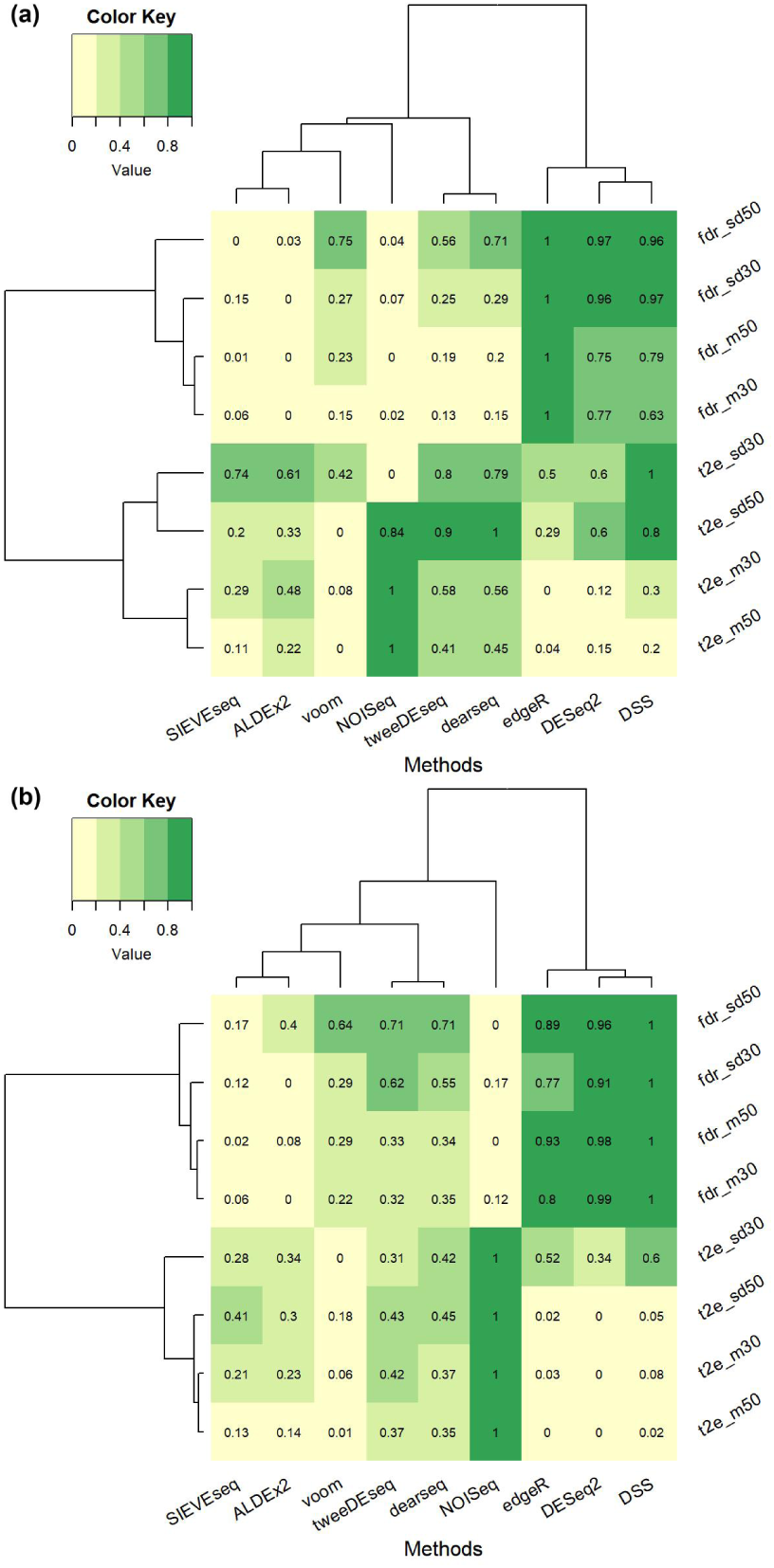
Cluster heatmaps for showing similarity of DE methods (columns) with respect to summary statistics (rows) of FDR and probability of Type II error (*β*) at two sample size scenarios (*n*=30, 50) for the (a) Dolgalev dataset and (b) Furusawa dataset. The number in a cell represents the mean (over 30 instances) of the summary statistic of a specific feature. Abbreviations: fdr = FDR; m = mean; sd = standard deviation; t2e = probability of Type II error; numerical suffixes indicate corresponding sample size scenario.

### 3.3 Analysis of the Mayo RNA-Seq Dataset

Most of the genes in both control (98.5%; 18385/18664) and AD groups (99.5%; 18571/18664) have CLR-transformed count data that are well-fitted (*p* -value >0.05) by the skew-normal distribution (Supplementary Fig. S5). We performed DE analysis of the Mayo RNA-Seq dataset by comparing SIEVEseq against edgeR, DESeq2, voom, tweeDEseq, ALDEx2, DSS, Wilcoxon, and NOISeq. We excluded dearseq because it is unable to provide output for the mean expression levels, rendering the significance score method used inapplicable. Supplementary Fig. S7 shows the number of DE genes unique to a DE method or common to multiple DE methods (for complete list, see Supplementary Table S13). DSS detected the most DE genes (4171), followed by DESeq2 (4155), edgeR (3942), voom (3636), tweeDEseq (3553), Wilcoxon (3330), ALDEx2 (3230), SIEVEseq (2773), and NOISeq (2014). That NOISeq detected the least number of DE genes is consistent with findings from the simulation studies that show its poor control of probability of Type II error. Supplementary Table S14 shows the list of DE, DV (2276) and DS (1024) genes detected using SIEVEseq (see also Supplementary Fig. S6). With respect to computational time, SIEVEseq completed the DE, DV, and DS tests concurrently in approximately 5.5 minutes. The run time for other methods (in increasing order) is as follows: 20s for DSS; about 30s for Wilcoxon; about 45s for edgeR, DESeq2, limma-voom and NOISeq; about 40 minutes for ALDEx2; and about 2.5 hours for tweeDEseq.

We defined the intersection of DE genes detected using each method as the high confidence gene set. Here, we excluded NOISeq because simulation studies suggested that it has a high false negative rate. Comparing SIEVEseq to three popular DE tests (i.e. edgeR, DESeq2 and voom), 98.7% (2736/2773) of DE genes identified by SIEVEseq are also detected by edgeR, DESeq2, or voom; 91.6% (2540/2773) are detected by edgeR, DESeq2, and voom; only 1.3% (37/2773) are uniquely detected by SIEVEseq. Furthermore, 93.1% (2583/2773), 94.3% (2615/2773) and 97.0% (2691/2773) of DE genes detected by SIEVEseq are also identified by edgeR, DESeq2 and voom, respectively. In comparison with ALDEx2, tweeDEseq, DSS and Wilcoxon, about 94.8% (2630/2733), 91.1% (2527/2773), 94.6% (2622/2773) and 91.2% (2528/2773) of DE genes identified by SIEVEseq are also detected by these four methods, respectively. Overall, 84.9% (2354/2773) of DE genes detected by SIEVEseq are also detected by all the other seven DE methods.

Table 3 gives the distribution of gene classes returned by SIEVEseq. The majority of genes are neither DE, DV, nor DS (∼ 70%; 13030/18664). For the remaining 30% of genes, focusing solely on DE genes means that only about 49% (2773/5634) of them would be considered in downstream analyses. The ability of SIEVEseq to detect the remaining 51% (2861/5634) of DV and/or DS genes that cannot be detected using DE methods sets it apart from all current methods. See Supplementary Fig. S8 to S12 for examples of distributional variation.

**Table 3.** Eight classes of genes identified by SIEVEseq for the Mayo RNA-Seq dataset. Abbreviations: 000 = non-DE, non-DV, and non-DS; 100 = DE, non-DV, and non-DS; 010 = non-DE, DV, and non-DS; 001 = non-DE, non-DV, and DS; 110 = DE, DV, and non-DS; 101 = DE, non-DV, and DS; 011 = non-DE, DV, and DS; 111 = DE, DV, and DS.

| Gene class | 000 | 100 | 010 | 001 | 110 | 101 | 011 | 111 |
| --- | --- | --- | --- | --- | --- | --- | --- | --- |
| Number of genes | 13030 | 2284 | 1843 | 919 | 303 | 155 | 99 | 31 |

Supplementary Table S7. summarizes the sets of pure DV, pure DS, and DVS genes detected using SIEVEseq that intersect with the DE gene sets identified by edgeR, DESeq2, voom, ALDEx2, tweeDEseq, DSS, and Wilcoxon. The key observation is that only 497/2861=17.4% of the genes in the union set of the pure DV, pure DS and DEV genes can be detected collaterally by the seven DE methods considered. This suggests that SIEVEseq detects a large number of genes with significant variability and skewness in gene expression distribution that standard DE methods fail to identify.

Figure 6 shows that DE, DV, and DS genes are represented among two alternative GUIDE trees for discriminating AD from the control state. The fibromodulin gene (FMOD; DE gene) is critical for enabling the identification of 70.7% (58/82) AD cases. A DS gene - tetratricopeptide repeat domain 7A (TTC7A), is crucial for enabling the prediction of the remaining AD cases (18.3% ;15/82) using the corticotropin releasing hormone gene (Fig. 8(a)). If TTC7A is removed, the resulting tree uses another DS gene - myotubularin related protein 7 gene (MTMR7) to predict the remaining 25.6% (21/82) AD cases (Fig. 8(b)). Overall, the first tree has estimated accuracy, sensitivity, and specificity of 87.5% (140/160), 89.0% (73/82) and 85.9% (67/78), respectively. For second tree, the estimated accuracy, sensitivity, and specificity is 89.4% (143/160), 96.3% (79/82) and 82.1% (64/78), respectively. See Supplementary Materials S5 for a brief summary of the biological relevance of genes in both trees.

**Fig. 8.**
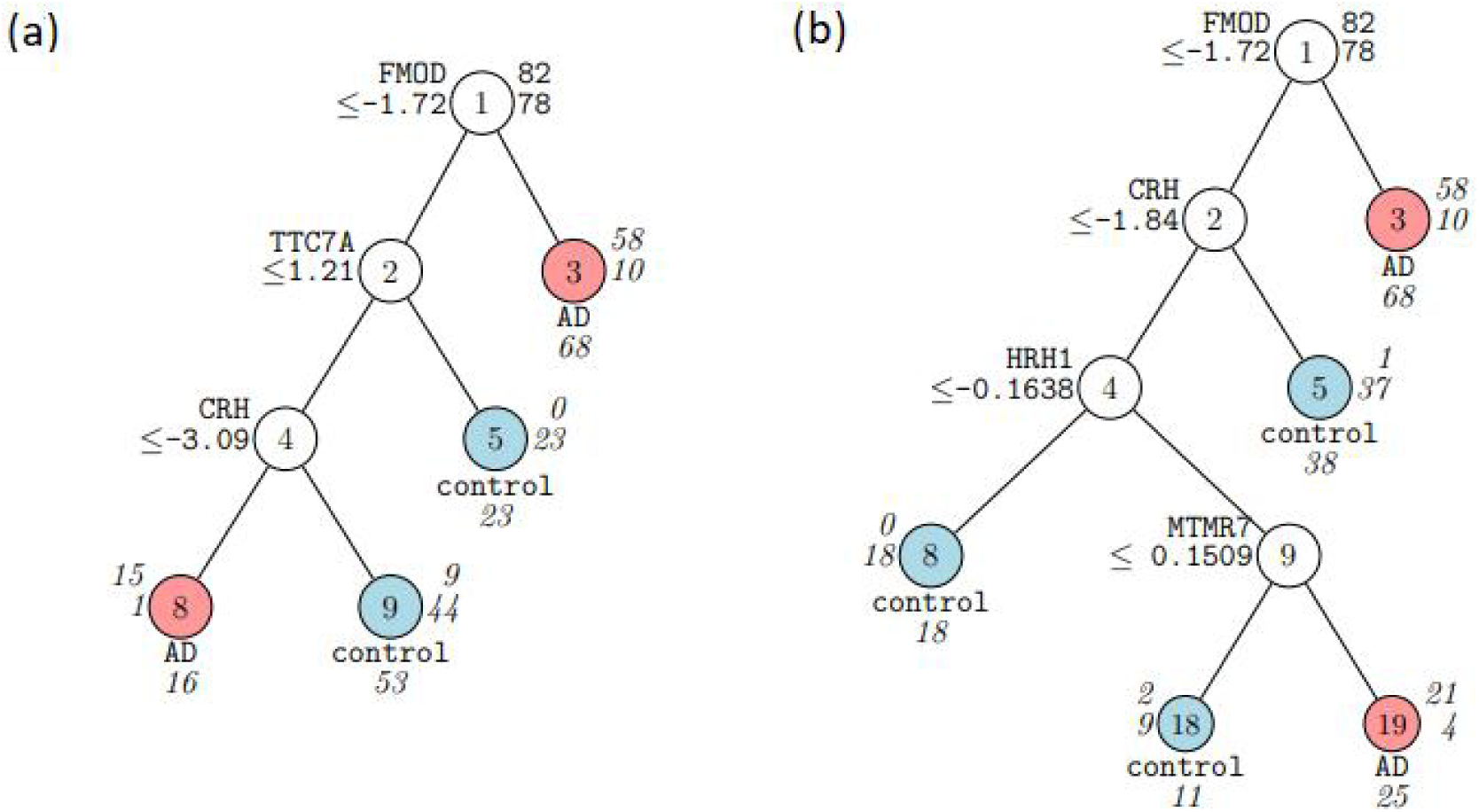
GUIDE v.40.3 0.250-SE classification tree for predicting using equal priors and unit misclassification costs. At each split, an observation goes to the left branch if and only if the condition is satisfied. Predicted classes and sample sizes (in *italics*) printed below terminal nodes; class sample sizes for = and beside nodes. (a) classification tree with *TTC7A*; (b) classification tree without *TTC7A*. *FMOD*: DE gene; *TTC7A*: DS gene; *CRH*: DE gene; *HRH1*: DV gene; *MTMR7*: DS gene.

The estimated generalization error of GUIDE was approximately the same (∼ 17%) regardless of whether DE, DV and DS gene sets were used separately, or collectively. In contrast, the estimated generalization error of GUIDE using genes that are neither DE, DV nor DS genes was more than twice as large (39.4%; 63/160). This finding suggests that the gene sets detected using SIEVEseq are informative.

The joint distribution of 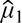 and 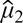 appears approximately bivariate normal (Fig. 9(a)), with DE genes distributed on the periphery of the outermost probability ellipse. Similarly, the joint distribution of log_2_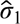 and log_2_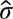_2_ is also approximately bivariate normal (Fig. 9(b)). Most genes have 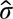 values between 0 and 1 in the control and the AD group, and are non-DV; DV genes generally have 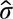 greater than 1, with most of them having smaller 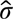 in the AD group compared to the control group (see also Li and Khang^30^). Finally, the joint distribution of 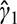 and 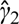 appears to be bimodal (Fig. 9(c)), with DS genes distributed towards the upper left and bottom right quadrants. In the AD group, genes tend to have 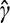 close to 0; thus, we may expect to see more genes with Gaussian-distributed CLR-transformed expression values. For genes in the control group, 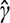 tends to be relatively more uniformly distributed; thus, distributions of CLR-transformed expression values are generally left or right-skewed.

**Fig. 9.**
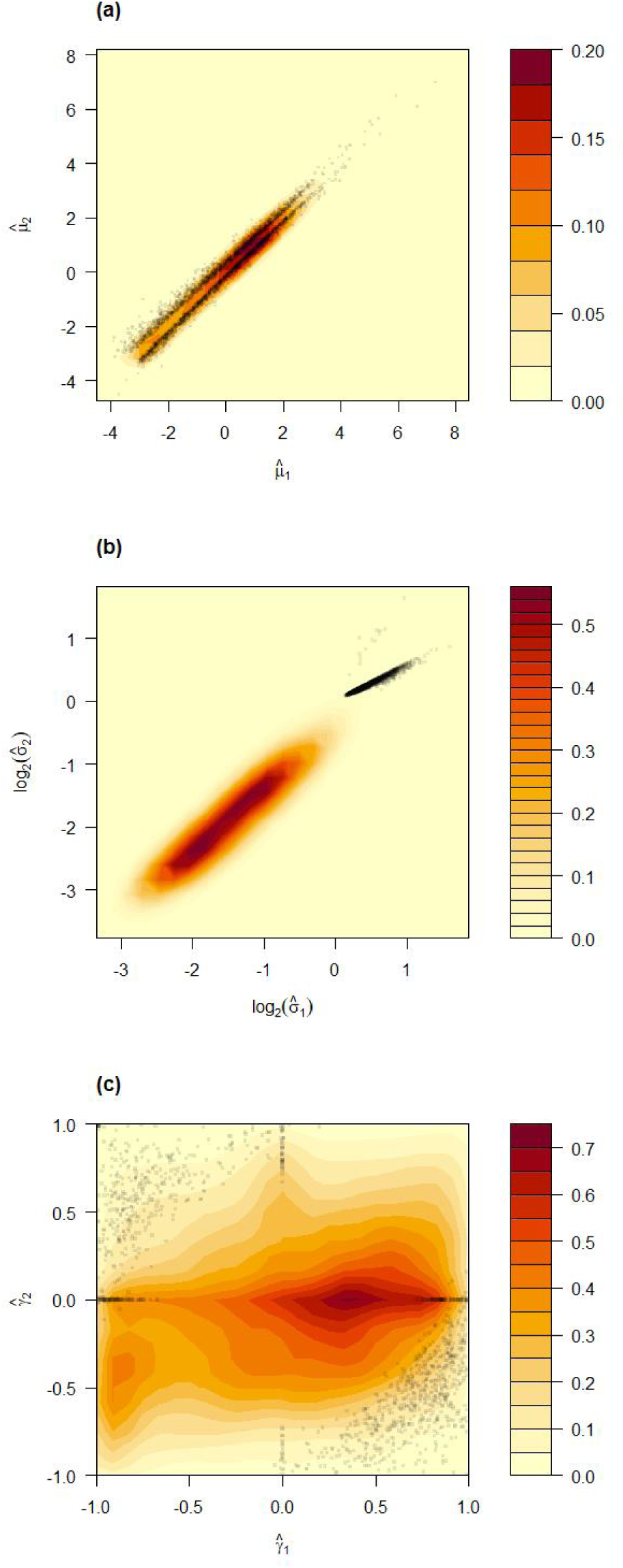
Contour plots of (a) 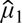 vs. 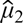; (b) log_2_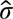_1_ vs. log2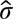_2_; and (c) 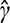_1_ vs. 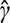_2_, with density color keys on the right. The subscripts 1 and 2 denote the control group and AD group, respectively. Significant genes are indicated as black squares. Range of the estimated parameters: 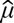_1_∈(-4.495, 8.457) ; log_2_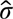_1_∈(-3.344, 1.867) ; 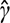_1_∈(-0.994, 0.994) ; 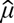_2_∈(-4.763, 8.190) ; log_2_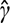_2_∈(-3.769, 1.817) ; 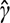_2_∈(-0.993, 0.994).

### 3.4 Functional Enrichment Analysis

Table 4 presents concise groupings of 18 GO terms obtained using all five gene lists into seven biological aspects. For details, see Supplementary Tables S8 to S12. The results indicate that an enrichment analysis that considers only conventional DE genes detects cell adhesion, extracellular structure organization, blood vessel development, and wound healing as enriched biological processes. Interestingly, consideration of non-DE but DV and/or DS genes reveals involvement of membrane protein localizations in AD pathogenesis, which is well-known, while simultaneously uncovering RNA catabolism. While the four enriched biological processes that are associated with pure DS gene list have FDR of about 0.61, they remain biologically interesting as studies of their involvement in AD have been reported. Importantly, enrichment analysis using the list of DE, DV, and DS genes from SIEVEseq not only recovers many hierarchically-related processes detected using just DE genes, or just DV and/or DS genes, but also uniquely detects the positive regulation of cell communication process. To summarize, our findings suggest that incorporating DV and DS analyses alongside DE analysis enables a more comprehensive understanding important biological processes involved in a disease.

**Table 4.** Functional enrichment in GO biological processes for the DE, DV, and DS genes for the control vs. AD comparison. The size of a gene list refers to the number of gene symbols in the list that are unambiguously mapped to unique Entrez gene IDs. For hierarchical GO terms, the reference GO term is indicated by [0]; parent/child terms have a +/- sign, with the numerical suffix indicating the hierarchy levels above/below the reference GO term. The FDRs of GO terms returned using all gene lists except that of pure DS have order of magnitude less than 10^-9^. For the latter gene list, the FDRs of the GO terms are all about 0.61. A: DE, DV, DS genes (4907); B: high confidence DE genes (1981); C: non-DE, DV, DS genes (2497); D: pure DV genes (1578); E: pure DS genes (811).

| Biological aspects | GO ID | GO description | Gene list (size) |  |  |  |  |
| --- | --- | --- | --- | --- | --- | --- | --- |
|  |  |  | A | B | C | D | E |
| Cellular protein localization | GO:0034613 | cellular protein localization [+4] |  |  | ✓ |  |  |
|  | GO:0072599 | establishment of protein localization to endoplasmic reticulum [+2] | ✓ |  |  |  |  |
|  | GO:0090150 | establishment of protein localization to membrane [+3] |  |  | ✓ | ✓ |  |
|  | GO:0045047 | protein targeting to ER [+1] |  |  |  | ✓ |  |
|  | GO:0006614 | SRP-dependent cotranslational protein targeting to membrane [0] |  |  | ✓ |  |  |
| Cell communication | GO:0022610 | biological adhesion [+2] | ✓ |  |  |  |  |
|  | GO:0031589 | cell-substrate adhesion [0] |  | ✓ |  |  |  |
|  | GO:0010647 | positive regulation of cell communication | ✓ |  |  |  |  |
|  | GO:0070848 | response to growth factor |  | ✓ |  |  |  |
| Matrix biology | GO:0030198 | extracellular matrix organization | ✓ |  |  |  |  |
| Angiogenesis | GO:0001568 | blood vessel development | ✓ | ✓ |  |  |  |
|  | GO:0042060 | wound healing |  | ✓ |  |  |  |
| Regulation of gene expression | GO:0006412 | translation |  |  | ✓ | ✓ |  |
|  | GO:0006401 | RNA catabolic process |  |  |  | ✓ |  |
| Immune response | GO:0019884 | antigen processing and presentation of exogenous antigen |  |  |  |  | ✓ |
|  | GO:0045619 | regulation of lymphocyte differentiation |  |  |  |  | ✓ |
| Oxidative stress | GO:0019363 | pyridine nucleotide biosynthetic process |  |  |  |  | ✓ |
|  | GO:0019752 | carboxylic acid metabolic process |  |  |  |  | ✓ |

A search of the AD literature reveals that many of these aspects in Table 4 have been investigated. Here, we briefly examine the literature that report on the association of these biological aspects with AD pathophysiology.

#### (i) Cellular protein localization

The endoplasmic reticulum (ER) is a critical organelle for the biosynthesis, folding, modification and assembly of proteins. Under stress, protein biosynthesis demand may exceed protein-folding capacity at the ER lumen, thus resulting in the production of proteins that are partially folded, misfolded, or unfolded, which leads to a condition known as ER stress.^62^ Amyloid beta (A*β*) deposition in the ER is thought to produce ER stress, and excessive levels of such stress can trigger maladaptive unfolded protein response (UPR) activation that produces excessive autophagy and induction of cell death pathways.^63^ Loss of the signal recognition particle (SRP) results in the mistargeting of genes bound for ER to the mitochondria, which damages mitochondria structure.^64^

#### (ii) Matrix biology

Constituents of the extracellular matrix (ECM) such as chondroitin sulfate proteoglycan and perineuronal net are potentially neuroprotective against tau lesions in AD pathogenesis.^65^ In particular, deglycosylation of perineuronal net has only very recently been shown to be associated with tauopathy-induced gliosis and neurodegeneration in mouse models.^66^

#### (iii) Cell communication

Cell adhesion molecules (CAMs) have been shown to be linked to A*β* metabolism as well as involved in neuroinflammatory responses and vascular changes.^67^ Dysfunctional components of Wnt signalling have been shown to be increase neuronal susceptibility to A *β* toxicity.^68^ Enhancing Wnt signalling may potentially mitigate synaptic pathology in AD.^69^

#### (iv) Angiogenesis

The involvement of angiogenesis in AD pathophysiology was hypothesized as early as the early 2000s^70^ and experimental support had accumulated since then.^71^ A fraction of AD cases may be caused by defects in brain wound healing.^72^ In a mouse model, overexpression of tau proteins induced changes in brain blood vessels that altered blood flow and led to cortical atrophy.^73^ Major findings implicating vascular channel dysfunction^74^ and pericyte-associated dysregulation of blood flow in brains of AD patients were reported in recent years.^75^

#### (v) Regulation of gene expression

The translation of the mRNA of the amyloid precursor protein (APP) is inhibited by the fragile X mental retardation protein (FMRP).^76,77^ Additionally, the reduced expression of an FMRP-binding protein known as the cytoplasmic FMRP-interacting protein 2 (CYFIP2) leads to accumulation of phosphorylated tau proteins and A *β* peptides.^78^ Meier et al. showed that tau-ribosome interaction in the ER impairs global RNA translation as well as the synthesis of an important synaptic protein that leads to synaptic dysfunction.^79^

#### (vi) Immune response

Neuroinflammation, whereby misfolded and aggregated proteins bind to surface receptors on microglia and astroglia to trigger innate immune response involving release of inflammatory mediators, is recognized as one of the hallmarks of AD pathophysiology.^80–82^ The effect of pathological differentiation of T cells during inflammation and the importance of antigen presentation through MHCII-positive microglia in AD pathophysiology were recently reviewed.^83,84^

#### (vii) Oxidative stress

The mitochondria, which is critical for cellular energy metabolism and redox homeostasis, is a major target of oxidative damage. Indeed, oxidative stress brought about by reactive oxygen species eventually leads to a runaway spiral towards cellular degeneration in AD.^85,86^ Pesini et al. hypothesized that the pathogenesis of late onset AD in a subgroup of AD patients may be related to a decrease in the de novo synthesis of pyrimidine nucleotides,^87^ which leads to dysfunction of the oxidative phosphorylation system. Carboxylic acid metabolism is related to lipid metabolism, and lipid dyshomeostasis has been shown to be present during early stages of AD brains.^88^

### 3.5 Cross-methodology and Cross-data Validations

Out of the 18 benchmark GO terms identified from the Mayo RNA-Seq dataset using ORA, 16 (∼89%) were successfully evaluated within the GSEA framework (Table 5). We found substantial functional consistency: about 37.5% (6/16) showed significant enrichment in GSEA, with adjusted *p* -value < 0.05. Relaxing the latter’s threshold to 0.20, about 56% (9/16) of the GO terms were significantly enriched. These substantial overlap percentages suggest that the identified biological processes reflect genuine biological signals and appear stable across different datasets. Such consistency confirms that the detected DE, DV and DS signals are likely robust and do not depend on a particular enrichment framework or a specific AD cohort. Thus, we conclude that inclusion of DV and DS genes enables the detection of signaling surveillance processes that govern heterogeneity of gene expression in cells, which is not possible with the use of DE genes only.

**Table 5.** Cross-methodology and cross-data validations of ORA-derived GO terms. Benchmark GO terms identified in the Mayo dataset were validated using GSEA in the independent Nakayama dataset. Enrichment was performed using a composite ranking score integrating DE, DV, and DS signals. NES = normalized enrichment score; adjusted p-value: < 0.05 (**), <0.20 (*).

| GO ID | Description | NES | Adjusted<br>*p*-value |
| --- | --- | --- | --- |
| GO:0010647 | positive regulation of cell communication | 1.49 | 0.0001** |
| GO:0045047 | protein targeting to ER | -1.85 | 0.0176** |
| GO:0019884 | antigen processing and presentation of exogenous antigen | 1.69 | 0.0268** |
| GO:0070848 | response to growth factor | 1.30 | 0.0268** |
| GO:0072599 | establishment of protein localization to endoplasmic reticulum | -1.69 | 0.0268** |
| GO:0045619 | regulation of lymphocyte differentiation | 1.40 | 0.0358** |
| GO:0006614 | SRP-dependent cotranslational protein targeting to membrane | -1.56 | 0.0891* |
| GO:0006401 | RNA catabolic process | 1.21 | 0.1190* |
| GO:0042060 | wound healing | 1.21 | 0.1190* |
| GO:0006412 | translation | -1.11 | 0.2766 |
| GO:0090150 | establishment of protein localization to membrane | -1.13 | 0.2772 |
| GO:0001568 | blood vessel development | 1.00 | 0.5868 |
| GO:0019752 | carboxylic acid metabolic process | -0.99 | 0.5868 |
| GO:0031589 | cell-substrate adhesion | 0.98 | 0.5868 |
| GO:0019363 | pyridine nucleotide biosynthetic process | 0.76 | 0.8491 |
| GO:0030198 | extracellular matrix organization | -0.64 | 0.9958 |

## 4. Discussion

In the Mayo RNA-Seq dataset, almost 50% (2773/5634) of genes that have significant difference in at least one aspect of the gene expression distribution can be detected using DE test. This affirms the value of conducting DE tests as a routine step in RNA-Seq data analysis. Importantly, DE tests cannot detect pure DV, pure DS, or DVS genes that constitute the remaining 50% (2861/5634) genes. As complex diseases are multifaceted, genomic studies that use DE genes for functional enrichment analysis would be disadvantaged by a narrow perspective. Here, querying the GO database using the gene lists generated by SIEVEseq substantially expands the landscape of biological processes that are potentially implicated in the pathogenesis of a disease, thus enabling their further exploration and investigation.

SIEVEseq may not be suitable for RNA-Seq studies with small sample sizes, since it generally requires larger sample sizes (50 or more per group) for reliable estimation of the standard deviation and skewness parameters. As the fit of the skew-normal model to CLR-transformed data is important, the result of testing of goodness-of-fit of the skew-normal model should be reported. Our present analysis was necessarily restricted to the Mayo RNA-Seq dataset, owing to the need for depth of analysis. We encourage more future analyses using SIEVEseq to ascertain whether the skew-normal distribution is indeed a ubiquitous feature of CLR-transformed RNA-Seq data.

To summarize, SIEVEseq unlocks the potential of RNA-Seq data by allowing a richer class of genes to be characterized with respect to their mean, standard deviation, and/or skewness characteristics. With the availability of SIEVEseq, substantial published RNA-Seq datasets of complex diseases may be fruitfully reanalyzed to generate new hypotheses concerning involvement of hitherto unconsidered biological processes. Another possible application is the annotation of genes in gene-gene interaction networks using the gene classes generated by SIEVEseq, which can potentially facilitate interpretation through integration of domain knowledge. Last but not least, given that pseudo-bulk profiles retain biological variability and skewness at the individual, cell-type, or group levels, SIEVEseq’s unique capacity to detect changes in these higher moments makes it a valuable tool for revealing important biological signals that are missed by conventional differential expression methods. This opens new opportunities for the reanalysis of large-scale pseudo-bulk scRNA-Seq datasets.

## Acknowledgements

The present work forms part of the PhD research of HL while at Universiti Malaya. Part of the manuscript was prepared while TFK was on sabbatical leave at the Institute for Chemical Research, Kyoto University. The results published here are in whole or in part based on data obtained from the AD Knowledge Portal. The Mayo RNAseq study data was led by Dr. Nilüfer Ertekin-Taner, Mayo Clinic, Jacksonville, FL as part of the multi-PI U01 AG046139 (MPIs Golde, Ertekin-Taner, Younkin, Price). Samples were provided from the following sources: The Mayo Clinic Brain Bank. Data collection was supported through funding by NIA grants P50 AG016574, R01 AG032990, U01 AG046139, R01 AG018023, U01 AG006576, U01 AG006786, R01 AG025711, R01 AG017216, R01 AG003949, NINDS grant R01 NS080820, CurePSP Foundation, and support from Mayo Foundation. Study data includes samples collected through the Sun Health Research Institute Brain and Body Donation Program of Sun City, Arizona. The Brain and Body Donation Program is supported by the National Institute of Neurological Disorders and Stroke (U24 NS072026 National Brain and Tissue Resource for Parkinsons Disease and Related Disorders), the National Institute on Aging (P30 AG19610 Arizona Alzheimers Disease Core Center), the Arizona Department of Health Services (contract 211002, Arizona Alzheimers Research Center), the Arizona Biomedical Research Commission (contracts 4001, 0011, 05-901 and 1001 to the Arizona Parkinson’s Disease Consortium) and the Michael J. Fox Foundation for Parkinsons Research. We would like to thank three anonymous reviewers and the handling editor Prof. Dr. Kenta Nakai for constructive comments which helped improve the present paper.

## References

1. Bahar, R., Hartmann, C.H., Rodriguez, K.A., et al. 2006, Increased cell-to-cell variation in gene expression in ageing mouse heart. Nature, 441, 1011–4.

2. Cheung, V.G., Conlin, L.K., Weber, T.M., et al. 2003, Natural variation in human gene expression assessed in lymphoblastoid cells. Nat Genet, 33, 422–5.

3. Pritchard, C.C., Hsu, L., Delrow, J., and Nelson, P.S. 2001, Project normal: Defining normal variance in mouse gene expression. PNAS, 98, 13266–71.

4. Ho, J.W.K., Stefani, M., Dos Remedios, C.G., and Charleston, M.A. 2008, Differential variability analysis of gene expression and its application to human diseases. Bioinformatics, 24, i390–8.

5. Mar, J.C., Matigian, N.A., Mackay-Sim, A., et al. 2011, Variance of gene expression identifies altered network constraints in neurological disease. PLoS Genet, 7, e1002207.

6. Gorlov, I.P., Byun, J., Zhao, H., Logothetis, C.J., and Gorlova, O.Y. 2012, Beyond comparing means: The usefulness of analyzing interindividual variation in gene expression for identifying genes associated with cancer development. J Bioinform Comput Biol, 10, 1241013.

7. Corrada Bravo, H., Pihur, V., McCall, M., Irizarry, R. A., and Leek, J.T. 2012, Gene expression anti-profiles as a basis for accurate universal cancer signatures. BMC Bioinformatics, 13, 272.

8. Casellas, J., and Varona, L. 2012, Modeling skewness in human transcriptomes. PLoS One, 7, e38919.

9. Marko, N.F., and Weil, R.J. 2012, Non-gaussian distributions affect identification of expression patterns, functional annotation, and prospective classification in human cancer genomes. PLoS One, 7, e46935.

10. Church, B., Williams, H.T., and Mar, J.C. 2019, Investigating skewness to understand gene expression heterogeneity in large patient cohorts. BMC Bioinformatics, 20, 668.

11. Mar, J.C. 2019, The rise of the distributions: Why non-normality is important for understanding the transcriptome and beyond. Biophys Rev, 11, 89–94.

12. Robinson, M.D., and Smyth, G.K. 2007, Moderated statistical tests for assessing differences in tag abundance. Bioinformatics, 23, 2881–7.

13. McCarthy, D.J., Chen, Y., and Smyth, G.K. 2012, Differential expression analysis of multifactor RNA-Seq experiments with respect to biological variation. Nucleic Acids Res, 40, 4288–97.

14. Love, M., Huber, W., and Anders, S. 2014, Moderated estimation of fold change and dispersion for RNA-seq data with DESeq2. Genome Biol, 15, 550.

15. Geiler-Samerotte, K.A., Bauer, C.R., Li, S., Ziv, N., Gresham, D., and Siegal, M.L. 2013, The details in the distributions: Why and how to study phenotypic variability. Curr Opin Biotechnol, 24, 752–9.

16. Srivastava, S., and Chen, L. 2010, A two-parameter generalized Poisson model to improve the analysis of RNA-seq data. Nucleic Acids Res, 38, e170.

17. Esnaola, M., Puig, P., Gonzalez, D., Castelo, R., and Gonzalez, J. R. 2013, A flexible count data model to fit the wide diversity of expression profiles arising from extensively replicated RNA-seq experiments. BMC Bioinformatics, 14, 254.

18. Law, C., Chen, Y., Shi, W., and Smyth, G.K. 2014, Voom: Precision weights unlock linear model analysis tools for RNA-seq read counts. Genome Biol, 15, R29.

19. Smyth, G.K. 2005, Limma: Linear models for microarray data Bioinformatics and Computational Biology Solutions Using R and Bioconductor. Springer, pp. 397–420.

20. Gauthier, M., Agniel, D., Thiébaut, R., and Hejblum, B.P. 2020, Dearseq: A variance component score test for RNA-seq differential analysis that effectively controls the false discovery rate. NAR Genom Bioinform, 2, lqaa093.

21. Tarazona, S., Furió-Tarí, P., Turrà, D., et al. 2015, Data quality aware analysis of differential expression in RNA-seq with NOISeq R/Bioc package. Nucleic Acids Res, 43, e140.

22. Tarazona, S., García-Alcalde, F., Dopazo, J., Ferrer, A., and Conesa, A. 2011, Differential expression in RNA-seq: A matter of depth. Genome Res, 21, 2213–23.

23. Li, J., and Tibshirani, R. 2013, Finding consistent patterns: A nonparametric approach for identifying differential expression in RNA-Seq data. Stat Methods Med Res, 22, 519–36.

24. Li, Y., Ge, X., Peng, F., Li, W., and Li, J.J. 2022, Exaggerated false positives by popular differential expression methods when analyzing human population samples. Genome Biol, 23, 79.

25. Phipson, B., and Oshlack, A. 2014, DiffVar: A new method for detecting differential variability with application to methylation in cancer and aging. Genome Biol, 15, 465.

26. Ran, D., and Daye, Z.J. 2017, Gene expression variability and the analysis of large-scale RNA-seq studies with the MDSeq. Nucleic Acids Res, 45, e127.

27. Rigby, R.A., and Stasinopoulos, D. M. 2005, Generalized additive models for location, scale and shape. J R Stat Soc Ser C Appl Stat, 54, 507–54.

28. de Jong, T.V., Moshkin, Y.M., and Guryev, V. 2019, Gene expression variability: The other dimension in transcriptome analysis. Physiol Genomics, 51, 145–58.

29. Roberts, A.G.K., Catchpoole, D.R., and Kennedy, P.J. 2022, Identification of differentially distributed gene expression and distinct sets of cancer-related genes identified by changes in mean and variability. NAR Genom Bioinform, 4, lqab124.

30. Li, H., and Khang, T.F. 2023, ClrDV: A differential variability test for RNA-Seq data based on the skew-normal distribution. PeerJ, 11, e16126.

31. Hasegawa, Y., Taylor, D., Ovchinnikov, D.A., Wolvetang, E.J., de Torrenté, L., and Mar, J.C. 2015, Variability of gene expression identifies transcriptional regulators of early human embryonic development. PLoS Genet, 11, e1005428.

32. Hardin, J., and Wilson, J. 2009, A note on oligonucleotide expression values not being normally distributed. Biostatistics, 10, 446–50.

33. Thomas, R., de la Torre, L., Chang, X., and Mehrotra, S. 2010, Validation and characterization of DNA microarray gene expression data distribution and associated moments. BMC Bioinformatics, 11, 576.

34. Fernandes, A.D., Macklaim, J.M., Linn, T.G., Reid, G., and Gloor, G.B. 2013, ANOVA-like differential expression (ALDEx) analysis for mixed population RNA-Seq. PLoS One, 8, e67019.

35. Fernandes, A.D., Reid, J.N., Macklaim, J.M., McMurrough, T.A., Edgell, D.R., and Gloor, G.B. 2014, Unifying the analysis of high-throughput sequencing datasets: Characterizing RNA-seq, 16S rRNA gene sequencing and selective growth experiments by compositional data analysis. Microbiome, 2, 15.

36. Quinn, T.P., Crowley, T.M., and Richardson, M.F. 2018, Benchmarking differential expression analysis tools for RNA-Seq: Normalization-based vs. Log-ratio transformation-based methods. BMC Bioinformatics, 19, 274.

37. Rasch, D., Teuscher, F., and Guiard, V. 2007, How robust are tests for two independent samples? J Stat Plan Inference, 137, 2706–20.

38. Aitchison, J. 1986, The Statistical Analysis of Compositional Data. Chapman & Hall, London.

39. Azzalini, A. 1985, A class of distributions which includes the normal ones. Scand J Stat, 12(2), 171–8.

40. Azzalini, A., and Capitanio, A. 2014, The Skew-Normal and Related Families. Cambridge University Press.

41. Azzalini, A., and Arellano-Valle, R.B. 2013, Maximum penalized likelihood estimation for skew-normal and skew-t distributions. J Stat Plan Inference, 143, 419–33.

42. Benjamini, Y., and Yekutieli, D. 2001, The control of the false discovery rate in multiple testing under dependency. Ann Stat, 29, 1165–88.

43. Leal Valentim, F., Mariotti-Ferrandiz, E., Klatzmann, D., Six, A., and Konza, O. 2020, Transimmunom whole blood RNA-seq data from type 1 diabetic patients and healthy volunteers.

44. Kelmer Sacramento, E., Kirkpatrick, J.M., Mazzetto, M., et al. 2020, Reduced proteasome activity in the aging brain results in ribosome stoichiometry loss and aggregation. Mol Syst Biol, 16, e9596.

45. Zhou, Y., Gallins, P.J., Etheridge, A.S., et al. 2022, A resource for integrated genomic analysis of the human liver. Sci Rep, 12, 15151.

46. Dolgalev I., Zhou H., Murrell N., et al. 2023, Inflammation in the tumor-adjacent lung as a predictor of clinical outcome in lung adenocarcinoma. Nat Commun, 14(1), 6764.

47. Furusawa H., Cardwell J.H., Okamoto T., et al. 2020, Chronic hypersensitivity pneumonitis, an interstitial lung disease with distinct molecular signatures. Am J Respir Crit Care Med. 202(10), 1430–44.

48. Allen, M., Carrasquillo, M.M., Funk, C., et al. 2016, Human whole genome genotype and transcriptome data for Alzheimer’s and other neurodegenerative diseases. Sci Data, 3, 160089. 10.1038/sdata.2016.89

49. Azzalini, A. 2022, The R package sn: The skew-normal and related distributions such as the skew-t and the SUN (version 2.1.0). Università degli Studi di Padova, Italia.Home page: Home page: http://azzalini.stat.unipd.it/SN/. Available from: https://cran.r-project.org/package=sn.

50. Iga, J.I., Yoshino, Y., Ozaki, T., et al. 2024, Blood RNA transcripts show changes in inflammation and lipid metabolism in Alzheimer’s disease and mitochondrial function in mild cognitive impairment. J Alzheimer’s Dis Rep, 8(1), 1690–1703.

51. Robinson, M.D., and Oshlack, A. 2010, A scaling normalization method for differential expression analysis of RNA-seq data. Genome Biol, 11, R25.

52. Frazee, A.C., Jaffe, A.E., Langmead, B., and Leek, J.T. 2015, Polyester: Simulating RNA-seq datasets with differential transcript expression. Bioinformatics, 31, 2778–84.

53. Benidt, S., and Nettleton, D. 2015, SimSeq: a nonparametric approach to simulation of RNA-sequence datasets. Bioinformatics, 31(13), 2131–40.

54. Xiao, Y., Hsiao, T.H., Suresh, U., et al. 2014, A novel significance score for gene selection and ranking. Bioinformatics, 30, 801–7.

55. Loh, W.Y. 2022, GUIDE (version 40.3). Available at: https://pages.cs.wisc.edu/loh/guide.html.

56. Loh, W.Y. 2009, Improving the precision of classification trees. Ann Appl Stat, 3, 1710–37.

57. Liao, Y., Wang, J., Jaehnig, E.J., Shi, Z., and Zhang, B. 2019, WebGestalt 2019: Gene set analysis toolkit with revamped UIs and APIs. Nucleic Acids Res, 47, W199–205.

58. Zhang, B., Kirov, S., and Snoddy, J. 2005, WebGestalt: An integrated system for exploring gene sets in various biological contexts. Nucleic Acids Res, 33, W741–8.

59. Subramanian, A., Tamayo, P., Mootha, V.K., et al. 2005, Gene set enrichment analysis: a knowledge-based approach for interpreting genome-wide expression profiles. PNAS, 102(43), 15545–50.

60. R Core Team. 2022, R: A Language and Environment for Statistical Computing. R Foundation for Statistical Computing, Vienna, Austria. Available from: https://www.r-project.org/.

61. Saurin, A. 2022, Bioinformatics tools for genomics and transcriptomics analyses: ENSEMBL ID to Gene Symbol Converter. Available at https://www.biotools.fr/human/ensembl_symbol_converter.

62. Schwarz, D.S., and Blower, M.D. 2016, The endoplasmic reticulum: Structure, function and response to cellular signaling. Cell Mol Life Sci, 73, 79–94.

63. Ajoolabady, A., Lindholm, D., Ren, J., and Pratico, D. 2022, ER stress and UPR in Alzheimer’s disease: Mechanisms, pathogenesis, treatments. Cell Death Dis, 13, 706.

64. Costa, E.A., Subramanian, K., Nunnari, J., and Weissman, J.S. 2018, Defining the physiological role of SRP in protein-targeting efficiency and specificity. Science, 359, 689–92.

65. Sethi, M.K., and Zaia, J. 2017, Extracellular matrix proteomics in schizophrenia and Alzheimer’s disease. Anal Bioanal Chem, 409, 379–94.

66. Logsdon, A.F., Foresi, B., Hu, S.J., et al. 2024, Perineuronal net deglycosylation associates with tauopathy-induced gliosis and neurodegeneration. J Neurochem, 168, 1923–36.

67. Wennström, M., and Nielsen, H.M. 2012, Cell adhesion molecules in Alzheimer’s disease. Degener Neurol Neuromuscul Dis, 2, 65–77.

68. Godoy, J.A., Rios, J.A., Zolezzi, J.M., Braidy, N., and Inestrosa, N.C. 2014, Signaling pathway cross talk in Alzheimer’s disease. J Cell Commun Signal, 12, 23.

69. Palomer, E., Buechler, J., and Salinas, P.C. 2019, Wnt signaling deregulation in the aging and Alzheimer’s brain. Front Cell Neurosci, 13, 227.

70. Vagnucci, A.H., and Li, W.W. 2003, Alzheimer’s disease and angiogenesis. Lancet, 361, 605–8.

71. Jefferies, W.A., Price, K.A., Biron, K.E., Fenninger, F., Pfeifer, C.G., and Dickstein, D.L. 2013, Adjusting the compass: New insights into the role of angiogenesis in Alzheimer’s disease. Alzheimers Res Ther, 5, 64.

72. Lehrer, S., and Rheinstein, P.H. 2016, A derangement of the brain wound healing process may cause some cases of Alzheimer’s disease. Discov Med, 22, 43.

73. Bennett, R.E., Robbins, A.B., Hu, M., et al. 2018, Tau induces blood vessel abnormalities and angiogenesis-related gene expression in P301L transgenic mice and human Alzheimer’s disease. PNAS, 115, E1289–98.

74. Taylor, J.L., Pritchard, H.A.T., Walsh, K.R., et al. 2022, Functionally linked potassium channel activity in cerebral endothelial and smooth muscle cells is compromised in Alzheimer’s disease. PNAS, 119, e2204581119.

75. Yang, A.C., Vest, R.T., Kern, F.N., et al. 2022, A human brain vascular atlas reveals diverse mediators of Alzheimer’s risk. Nature, 603, 885–92.

76. Westmark, C.J., and Malter, J.S. 2007, FMRP mediates mGluR5-dependent translation of amyloid precursor protein. PLoS Biol, 5, e52.

77. Li, Z., Zhang, Y., Ku, L., Wilkinson, K.D., Warren, S.T., and Feng, Y. 2001, The fragile X mental retardation protein inhibits translation via interacting with mRNA. Nucleic Acids Res, 29, 2276–83.

78. Ghosh, A., Mizuno, K., Tiwari, S.S., et al. 2020, Alzheimer’s disease-related dysregulation of mRNA translation causes key pathological features with ageing. Transl Psychiatry, 10, 192.

79. Meier, S., Bell, M., Lyons, D.N., et al. 2016, Pathological tau promotes neuronal damage by impairing ribosomal function and decreasing protein synthesis. J Neurosci, 36, 1001–7.

80. Akiyama, H., Barger, S., Barnum, S., et al. 2000, Inflammation and Alzheimer’s disease. Neurobiol Aging, 21, 383–421.

81. Cameron, B., and Landreth, G.E. 2010, Inflammation, microglia, and Alzheimer’s disease. Neurobiol Dis, 37, 503–9.

82. Heneka, M.T., Carson, M.J., El Khoury, J., et al. 2015, Neuroinflammation in Alzheimer’s disease. Lancet Neurol, 14, 388–405.

83. Schetters, S.T., Gomez-Nicola, D., Garcia-Vallejo, J.J., and Van Kooyk, Y. 2018, Neuroinflammation: Microglia and T cells get ready to tango. Front Immunol, 8, 1905.

84. Moro-García, M.A., Mayo, J. C., Sainz, R. M., and Alonso-Arias, R. 2018, Influence of inflammation in the process of T lymphocyte differentiation: Proliferative, metabolic, and oxidative changes. Front Immunol, 9, 328039.

85. Wang, X., Wang, W., Li, L., Perry, G., Lee, H., and Zhu, X. 2014, Oxidative stress and mitochondrial dysfunction in Alzheimer’s disease. Biochim Biophys Acta, 1842, 1240–7.

86. Swerdlow, R.H., Burns, J.M., and Khan, S.M. 2014, The Alzheimer’s disease mitochondrial cascade hypothesis: Progress and perspectives. Biochim Biophys Acta, 1842, 1219–31.

87. Pesini, A., Iglesias, E., Bayona-Bafaluy, M.P., et al. 2019, Brain pyrimidine nucleotide synthesis and Alzheimer disease. Aging, 11, 8433.

88. Yin, F. 2023, Lipid metabolism and Alzheimer’s disease: Clinical evidence, mechanistic link and therapeutic promise. FEBS J, 290, 1420–53.

