## Supplementary Materials for "SIEVEseq: One-stop differential expression, variability, and skewness analyses using RNA-Seq data"

#### S1 Statistical Modeling of CLR-transformed RNA-Seq Data

##### S1.1 The Skew-normal Distribution

Let $\phi(x)$ be the standard normal probability density function (pdf) $\phi(x) = \frac{1}{\sqrt{2\pi}}e^{-\frac{x^2}{2}}$, with cumulative distribution function $\Phi(x) = \int_{-\infty}^{x} \phi(t)\, dt$. Suppose $Z$ is a skew-normal random variable. Its probability density function is given by twice of the product of $\phi(z)$ and $\Phi(\alpha z)$:

$$\varphi(z, \alpha) = 2\phi(z)\Phi(\alpha z),$$

where $\alpha \in \mathbb{R}$ is the skewness parameter. We shall denote a skew-normal random variable with skewness parameter $\alpha$ as $Z \sim \mathrm{SN}(\alpha)$.

The mean and the variance of $Z$ are given by $\mu_z = \mathbb{E}(Z) = b\delta$, and $\sigma_z^2 = \mathrm{Var}(Z) = 1 - (b\delta)^2$, respectively, where $b = \sqrt{2/\pi}$ and $\delta = \alpha/\sqrt{1+\alpha^2}$. The formal derivation of the properties of the skew-normal distribution is due to Azzalini (1985), who treated the skew-normal distribution as a generalization of the normal distribution. Different authors derived the skew-normal model in other contexts (e.g. as a prior distribution in Bayesian analysis by O'Hagan and Leonard (1976); see Azzalini (2022)), but did not follow up with the extensive elaborations as done by Azzalini (1985).

The random variable $Y = \xi + \omega Z$ is a skew-normal variable with location parameter $\xi$, scale parameter $\omega$ and skewness parameter $\alpha$, where $\xi \in \mathbb{R}$ and $\omega \in \mathbb{R}^+$. We shall denote it as $Y \sim \mathrm{SN}(\xi, \omega, \alpha)$. Its pdf is given by

$$f(y;\ \xi, \omega, \alpha) = \frac{2}{\omega}\phi\Big(\frac{y-\xi}{\omega}\Big)\Phi\Big(\alpha\frac{y-\xi}{\omega}\Big).$$

We say that $Y$ follows a skew-normal distribution with direct parameters (DP) $\boldsymbol{\theta}^{(D)} = (\xi, \omega, \alpha)$. The mean, the variance, and the skewness of $Y$ are given by

$$\mathrm{E}(Y) = \xi + \omega\mu_z, \tag{1}$$

$$\mathrm{Var}(Y) = (\omega\sigma_z)^2, \tag{2}$$

$$\gamma(Y) = \left(2 - \frac{\pi}{2}\right)\frac{\mu_z^3}{\sigma_z^3},$$

respectively. Azzalini (1985) found that the Fisher information matrix for $\boldsymbol{\theta}^{(D)}$ becomes singular as $\alpha \to 0$. To avoid this singularity, he redefined $Y$ as

$$Y = \mu + \sigma\frac{Z - \mu_z}{\sigma_z},$$

where $\mu = \mathrm{E}(Y)$ is given by Eq. (1), $\sigma = \mathrm{Var}(Y)$ is given by Eq. (2), and the skewness parameter $\gamma$ is the coefficient of skewness of $Z$, and also that of $Y$. The centered parameters (CP) vector $\boldsymbol{\theta}^{(C)} = (\mu, \sigma, \gamma)$ has parameter space $\mathbb{R} \times \mathbb{R}^+ \times (-k, k)$, where $k = \sqrt{2}(4-\pi)/(\pi-2)^{3/2} \approx 0.9953$. We shall write $Y \sim \mathrm{SN}_C(\mu, \sigma, \gamma)$ to indicate a skew-normal random variable with centered parameters. The special case of $\gamma = 0$ reduces to the normal distribution with mean $\mu$ and variance $\sigma^2$ (Azzalini, 1985; Azzalini and Capitanio, 2014). The skew-normal distribution with CP is derived from the DP form via the mapping (Azzalini and Capitanio, 2014)

$$\begin{aligned} \mu_g &= \xi_g + b\omega_g\delta_g, \\ \sigma_g &= \omega_g\sqrt{1 - b^2\delta_g^2}, \\ \gamma_g &= \frac{4-\pi}{2}\frac{b^3\alpha_g^3}{\left\{1 + (1-b^2)\alpha_g^2\right\}^{3/2}}; \end{aligned} \tag{3}$$

and the inverse mapping is provided by

$$\begin{aligned} \xi_g &= \mu_g - b\omega_g\delta_g, \\ \omega_g &= \frac{\sigma_g}{\sqrt{1 - b^2\sigma_g^2}}, \\ \alpha_g &= \frac{R}{\sqrt{b^2 - (1-b^2)R^2}}, \end{aligned} \tag{4}$$

where $b = \sqrt{2/\pi}$, $\delta_g = \alpha_g/\sqrt{1+\alpha_g^2}$, and $R = \sqrt[3]{2\gamma_g/(4-\pi)}$.

#### S1.2 Parameter Estimation

For a single sample, the log-likelihood function for $\boldsymbol{\theta}_g^{(D)} = (\xi_g, \omega_g, \alpha_g)^T$ is given by

$$\ell_1 = \log L(\boldsymbol{\theta}_g^{(D)};\ y_{gi}) = c - \log\omega_g - \frac{(y_{gi} - \xi_g)^2}{2\omega_g^2} + \zeta_0\Big(\alpha_g\frac{y_{gi} - \xi_g}{\omega_g}\Big),$$

where $c$ is a constant and $\zeta_0(\cdot) = \log\{2\Phi(\cdot)\}$. Taking

$$z_{gi} = \frac{y_{gi} - \xi_g}{\omega_g},$$

we obtain the partial derivatives of $\ell_1$:

$$\frac{\partial\ell_1}{\partial\xi_g} = \frac{z_{gi}}{\omega_g} - \frac{\alpha_g}{\omega_g}\zeta_1(\alpha_g z_{gi}),$$

$$\frac{\partial \ell_1}{\partial \omega_g} = -\frac{1}{\omega_g} + \frac{z_{gi}^2}{\omega_g} - \frac{\alpha_g}{\omega_g}\zeta_1(\alpha_g z_{gi}) z_{gi},$$
$$\frac{\partial \ell_1}{\partial \alpha_g} = \zeta_1(\alpha_g z_{gi}) z_{gi};$$

thus the likelihood equations for a sample of size $n$ are given by

$$\begin{aligned}
&\sum_{i=1}^{n} z_{gi} - \alpha_g \sum_{i=1}^{n} \zeta_1(\alpha_g z_{gi}) = 0, \\
&\sum_{i=1}^{n} {z_{gi}}^2 - \alpha_g \sum_{i=1}^{n} z_{gi}\zeta_1(\alpha_g z_{gi}) = n, \\
&\sum_{i=1}^{n} z_{gi}\zeta_1(\alpha_g z_{gi}) = 0,
\end{aligned} \tag{5}$$

where $\zeta_1(\cdot) = \phi(\cdot)/\Phi(\cdot)$. The solution of these equations requires numerical methods because of the presence of $\zeta_1$, which is a non-linear function. Additionally, sample sizes of 50 or more are recommended (Azzalini and Capitanio, 2014). To initialize the search, method of moments (MM) estimates are chosen as starting points for the CP components in Eq. (3). The MM estimators for the centered parameters are given by

$$\tilde{\mu}_g = \bar{Y}_g, \quad \tilde{\sigma}_g = s_g, \quad \tilde{\gamma}_g = \frac{M_{g,3}}{s_g^3}, \tag{6}$$

respectively, where $\bar{Y}_g$ is the sample mean, $s_g$ is the sample standard deviation, and $M_{g,3}$ is the sample third central moment. By estimating the CP components in Eq. (3) using Eq. (6), and then converting them to DP components using Eq. (4), we obtain the MM estimators of the DP components: $\bar{\xi}_g$, $\bar{\omega}_g$ and $\bar{\alpha}_g$. Subsequently, a search of the DP space where Eq. (5) holds is done. Once $\hat{\boldsymbol{\theta}}_g^{(D)} = (\hat{\xi}_g, \hat{\omega}_g, \hat{\alpha}_g)$ is obtained, it is mapped to Eq. (3) to obtain $\hat{\boldsymbol{\theta}}_g^{(C)} = (\hat{\mu}_g, \hat{\sigma}_g, \hat{\gamma}_g)$, the maximum likelihood estimators of the centered parameters.

#### S1.3 Fisher Information Matrix

For the DP vector $\boldsymbol{\theta}^{(D)} = (\xi, \omega, \alpha)$, the Fisher information matrix is given by

$$\boldsymbol{I}_{\boldsymbol{\theta}^{(D)}} = \begin{bmatrix} \frac{1+\alpha^2 a_0}{\omega^2} & \frac{1}{\omega^2}\Big(\mathrm{E}(Z)\frac{1+2\alpha^2}{1+\alpha^2} + \alpha^2 a_1\Big) & \frac{1}{\omega}\Big\{\frac{b}{(1+\alpha^2)^{3/2}} - \alpha a_1\Big\} \\ \frac{1}{\omega^2}\Big(\mathrm{E}(Z)\frac{1+2\alpha^2}{1+\alpha^2} + \alpha^2 a_1\Big) & \frac{2+\alpha^2 a_1}{\omega^2} & -\frac{\alpha a_2}{\omega} \\ \frac{1}{\omega}\Big\{\frac{b}{(1+\alpha^2)^{3/2}} - \alpha a_1\Big\} & -\frac{\alpha a_2}{\omega} & a_2 \end{bmatrix},$$

where $b = \sqrt{2/\pi}$, $a_k = a_k(\alpha) = \mathrm{E}\Big(Z^k \zeta_1^2(\alpha Z)\Big), \zeta_1(\cdot) = \phi(\cdot)/\Phi(\cdot), k = 0, 1, 2$ (Azzalini, 1985). For the CP vector $\boldsymbol{\theta}^{(C)} = (\mu, \sigma, \gamma)$, the Fisher information matrix can be found from $\boldsymbol{I}_{\boldsymbol{\theta}^{(D)}}$ as

$$\boldsymbol{I}_{\boldsymbol{\theta}^{(C)}} = \boldsymbol{D}^T \boldsymbol{I}_{\boldsymbol{\theta}^{(D)}} \boldsymbol{D},$$

where $\boldsymbol{D}$ is the matrix of the derivatives of the parameters $\boldsymbol{\theta}^{(D)} = (\xi, \omega, \alpha)$ (i.e. the Jacobian matrix) with respect to $\boldsymbol{\theta}^{(C)} = (\mu, \sigma, \gamma)$. We have

$$\boldsymbol{D} = \begin{bmatrix} 1 & -\frac{\mu_z}{\sigma_z} & \frac{\partial}{\partial\gamma}\xi \\ 0 & \frac{1}{\sigma_z} & \frac{\partial}{\partial\gamma}\omega \\ 0 & 0 & \frac{\partial}{\partial\gamma}\alpha \end{bmatrix},$$

where $\mu_z = b\delta$ and $\sigma_z^2 = 1 - b^2\delta^2$. The third column vector of $\boldsymbol{D}$ consists of

$$\frac{\partial}{\partial\gamma}\xi = -\frac{\sigma\mu_z}{3\sigma_z\gamma}, \quad \frac{\partial}{\partial\gamma}\omega = -\frac{\sigma}{\sigma_z^2}\frac{d\sigma_z}{d\alpha}\frac{d\alpha}{d\gamma}, \quad \frac{\partial}{\partial\gamma}\alpha = \frac{2}{3(4-\pi)}\Big(\frac{1}{TR^2} + \frac{1-b^2}{T^3}\Big),$$

where

$$\frac{d\sigma_z}{d\alpha} = -\frac{\mu_z}{\sigma_z}\frac{b}{(1+\alpha^2)^{3/2}}, \quad T = \sqrt{b^2 - (1-b^2)R^2}, \quad R = \left(\frac{2\gamma}{4-\pi}\right)^{1/3}.$$

The Fisher information matrix $\boldsymbol{I}_{\boldsymbol{\theta}^{(C)}}$ converges to a diagonal matrix with diagonal entries $(1/\sigma^2, 2/\sigma^2, 1/6)$, as $\gamma \to 0$ (Azzalini, 1985; Azzalini and Capitanio, 2014).

#### S1.4 Maximum Penalized Likelihood Estimation

Certain data values can result in a divergent $\hat{\alpha}_g$ for regular maximum likelihood estimation. To circumvent this problem, Azzalini and Arellano-Valle (2013) proposed a maximum penalized likelihood estimation ("Qpenalty") approach. We write the penalized log-likelihood for $\boldsymbol{\theta}_g^D = (\xi_g, \omega_g, \alpha_g)$ as

$$\ell_p(\boldsymbol{\theta}_g^{(D)}) = \ell(\boldsymbol{\theta}_g^{(D)};\ \boldsymbol{y}_g) - Q,$$

where $\boldsymbol{y}_{gi} = (y_{g1}, y_{g2}, \ldots, y_{gn})$, $\ell(\boldsymbol{\theta}_g^{(D)};\ \boldsymbol{y}_g)$ is the log-likelihood function with respect to the set to direct parameters $\boldsymbol{\theta}_g^D = (\xi_g, \omega_g, \alpha_g)$. The penalty function $Q$ satisfies

$$Q \geq 0, \quad Q|_{\alpha_g=0} = 0, \quad \lim_{|\alpha_g|\to\infty} Q = +\infty,$$

and $Q$ does not depend on $n$. The $Q$ penalty is formulated as

$$Q = c_1 \log(1 + c_2\alpha_g^2), \tag{7}$$

where $c_1 \approx 0.87591$ and $c_2 \approx 0.85625$ (Azzalini and Arellano-Valle, 2013; Azzalini and Capitanio, 2014). The maximum penalized likelihood estimator (MPLE), $\tilde{\boldsymbol{\theta}}_g^{(D)}$, maximizes $\ell_p(\boldsymbol{\theta}_g^{(D)})$. The standard errors of $\tilde{\boldsymbol{\theta}}_g^{(D)}$ can be approximated using the corresponding penalized information matrix as

$$\text{Var}(\tilde{\boldsymbol{\theta}}_g^{(D)}) \approx -\ell_p^{''}(\tilde{\boldsymbol{\theta}}_g^{(D)})^{-1}.$$

For the "MPpenalty" approach (Azzalini and Capitanio, 2014), the penalty function $Q$ in Eq. (7) is defined as $Q = -\log \pi_m(\alpha_g)$, where $\pi_m$ is a prior distribution for the skewness parameter $\alpha_g$. The matching prior (Cabras et al, 2012) for $\alpha_g$, allowing for the presence of $\boldsymbol{\psi} = (\xi_g, \omega_g)$, is given by

$$\pi_m(\alpha_g) \propto \left(I_{\alpha_g\alpha_g}(\hat{\boldsymbol{\psi}}, \alpha_g) - I_{\alpha_g\boldsymbol{\psi}}(\hat{\boldsymbol{\psi}}, \alpha_g) I_{\boldsymbol{\psi}\boldsymbol{\psi}}(\hat{\boldsymbol{\psi}}, \alpha_g)^{-1} I_{\boldsymbol{\psi}\alpha_g}(\hat{\boldsymbol{\psi}}, \alpha_g)\right)^{1/2},$$

where the terms involved are specific blocks of the Fisher information matrix $\boldsymbol{I}$ of $\boldsymbol{\theta}_g^{(D)}$. Since $\pi_m(0) = 0$, the matching prior penalty effectively penalizes $\alpha_g = 0$ with $Q = \infty$.

#### S1.5 Performance Evaluation Metrics

To evaluate the comparative performance of SIEVEseq and existing DE methods, we used several statistical metrics. Let $TP$, $FP$, $TN$, and $FN$ represent the number of true positives, false positives, true negatives, and false negatives, respectively, derived from the test results of a single simulated dataset. The metrics are defined as follows:

- **False Discovery Proportion (FDP):** The proportion of falsely identified DE genes among all genes detected significant, defined as:

$$FDP = \begin{cases} \frac{FP}{TP+FP}, & \text{if } TP + FP > 0 \\ 0, & \text{if } TP + FP = 0. \end{cases}$$

- **False Discovery Rate (FDR):** The $FDR$ was estimated by averaging the $FDP$ values across all independent simulation instances for a given scenario:

$$FDR = E[FDP] \approx \frac{1}{M} \sum_{m=1}^{M} FDP_m,$$

where $M$ is the total number of simulation runs (e.g., $M = 30$).

- **Probability of Type II Error ($\beta$):** Also known as the false negative rate, $\beta$ represents the probability of failing to reject a false null hypothesis (i.e., failing to detect a true DE gene):

$$\beta = \frac{FN}{TP + FN}.$$

- **Desirable performance region index ($\theta$):** This composite metric quantifies the reliability of a method by calculating the percentage of simulated instances that satisfy chosen performance criteria for FDR $< c_1$ and $\beta < c_2$:

$$\theta = \frac{1}{M} \sum_{m=1}^{M} I(FDR < c_1 \text{ and } \beta < c_2) \times 100\%,$$

where $I(\cdot)$ is the indicator function. In our study, $c_1$ and $c_2$ were set to 0.05 and 0.05 (or 0.2), respectively.

#### S1.6 Computational Aspects

To perform parameter estimation and carry out related numerical tasks involving the skew-normal distribution, we used the `sn` (Azzalini, 2022) R package. Regular maximum likelihood estimation of parameters of the skew-normal model was first done using the function `selm()`. If NA values were returned, the maximum penalized likelihood estimation as implemented using the `Qpenalty` option was used. If NA values persisted, the `MPpenalty` option was then used. Thus, numerically stable parameter estimates were obtained.

### S2 The Significance Score Method

Xiao et al (2014) proposed the significance score method which jointly considers statistical and biological significance when deciding whether a gene should be called as differentially expressed. The method specifies a significance score ($\pi$), which is the product of a measure of biological significance ($\eta$, the $\log_2$ fold-change or the difference between the mean of gene expression between two groups) and a measure of the desired level of statistical significance ($-\log_{10} p$, where $p$ is the $p$-value of the statistical test used, adjusted for multiple comparisons). Decision boundaries may be specified using desirable $\eta$ values (e.g. a lower value $\eta_l$ for $\eta < 0$ and an upper value $\eta_u$ for $\eta > 0$, where such values may be set by the user or determined from quantiles of estimated $\eta$ values over all genes tested) and a desired level of statistical significance $p'$. Thus, genes with $\pi > \eta_u(-\log_{10} p')$ or $\pi < \eta_l(-\log_{10} p')$ are flagged as DE genes. Although the method was originally designed for calling DE genes, it can also be used for calling DV and DS genes in the present case by choosing a suitable measure of $\eta$.

For all methods, we defined the threshold for statistical significance as an adjusted $p$-value of 0.05 or less. Since the CLR-transformed count data in SIEVEseq and ALDEx2 is a real number, we measured biological significance as the difference between mean CLR-transformed counts between two groups ($\Delta\hat{\mu}_g = \hat{\mu}_{g,2} - \hat{\mu}_{g,1}$). We specified the lower and the upper thresholds for biological significance as the 0.025 and the 0.975 quantiles of $\Delta\hat{\mu}_g$ of all genes tested for DE. Thus, genes with significance score of $\pi_g < (\Delta\hat{\mu}_g)_{0.025} \times (-\log_{10} 0.05)$ or $\pi_g > (\Delta\hat{\mu}_g)_{0.975} \times (-\log_{10} 0.05)$ were flagged as DE genes. For count-based methods (edgeR, DESeq2, voom, tweeDEseq and DSS), the lower and upper thresholds for biological significance were set as the 0.025 and the 0.975 quantiles of the $\log_2$ fold-change (i.e. $\log_2(\hat{\mu}_{g,2}/\hat{\mu}_{g,1})$).

For DV test, we defined biological significance as $\hat{\psi}_g = \log_2(\hat{\sigma}_{g,2}/\hat{\sigma}_{g,1})$ (i.e. $\log_2$ fold-change of SD ratio). We specified $|\hat{\psi}_g| > 1.25$ for biological significance. Thus, genes with significance score of

$$|\pi_g| = |\hat{\psi}_g| \times (-\log_{10} 0.05) > 1.25 \times (-\log_{10} 0.05)$$

were flagged as DV genes. Finally, for DS test, we defined biological significance as $\Delta\hat{\gamma}_g = \hat{\gamma}_{g,2} - \hat{\gamma}_{g,1}$, which is the difference between skewness of the distribution of CLR-transformed counts between two groups. We specified the lower and the upper thresholds for biological significance as the 0.025 and the 0.975 quantiles of $\Delta\hat{\gamma}_g$ of all genes tested for DS. Thus, genes with significance scores of $\pi_g < (\Delta\hat{\gamma}_g)_{0.025} \times (-\log_{10} 0.05)$ or $\pi_g > (\Delta\hat{\gamma}_g)_{0.975} \times (-\log_{10} 0.05)$ were flagged as DS genes.

### S3 Supplementary Tables

**Table S1** The mean of false discovery rate (FDR*), mean probability of Type II error ($\beta^{\dagger}$), and percentage of simulated instances with FDR < 0.05 and $\beta < 0.05$ ($\theta^{\ddagger}$) for each of the ten DE tests applied to data simulated from the Kelmer dataset (30 instances) at three different sample sizes. Standard deviation in parentheses.

| Method | Sample size per group | | |
|---|---|---|---|
| | 30 | 50 | 100 |
| SIEVEseq | 0.016 (0.009) | 0.013 (0.007) | 0.013 (0.009)* |
| | 0.005 (0.004) | 0.003 (0.004) | 0.001 (0.002)† |
| | 100% | 100% | 100%‡ |
| ALDEx2 | 0.030 (0.011) | 0.033 (0.013) | 0.035 (0.018) |
| | 0.011 (0.008) | 0.005 (0.006) | 0.002 (0.003) |
| | 93.3% | 93.3% | 80% |
| NOISeq | 0.002 (0.004) | 0.001 (0.002) | 0.001 (0.003) |
| | 0.049 (0.019) | 0.012 (0.008) | 0.000 (0.001) |
| | 53.3% | 100% | 100% |
| edgeR | 0.041 (0.013) | 0.038 (0.016) | 0.042 (0.016) |
| | 0.001 (0.002) | 0.000 (0.000) | 0.000 (0.000) |
| | 83.3% | 83.3% | 70% |
| DESeq2 | 0.056 (0.017) | 0.051 (0.017) | 0.049 (0.014) |
| | 0.001 (0.002) | 0.000 (0.001) | 0.000 (0.000) |
| | 43.3% | 46.7% | 43.3% |
| voom | 0.042 (0.012) | 0.045 (0.019) | 0.046 (0.019) |
| | 0.003 (0.003) | 0.002 (0.003) | 0.001 (0.002) |
| | 76.7% | 70% | 56.7% |
| DSS | 0.074 (0.020) | 0.065 (0.016) | 0.055 (0.014) |
| | 0.001 (0.002) | 0.000 (0.001) | 0.000 (0.000) |
| | 6.7% | 23.3% | 26.7% |
| dearseq | 0.061 (0.016) | 0.049 (0.016) | 0.052 (0.015) |
| | 0.001 (0.003) | 0.000 (0.000) | 0.000 (0.000) |
| | 26.7% | 56.7% | 43.3% |
| Wilcoxon | 0.042 (0.015) | 0.042 (0.017) | 0.044 (0.017) |
| | 0.002 (0.003) | 0.000 (0.001) | 0.000 (0.000) |
| | 76.7% | 73.3% | 67.7% |
| tweeDEseq | 0.056 (0.016) | 0.047 (0.016) | 0.049 (0.015) |
| | 0.002 (0.003) | 0.000 (0.000) | 0.000 (0.000) |
| | 36.7% | 53.3% | 46.7% |

**Table S2** The mean of false discovery rate (FDR*), mean probability of Type II error ($\beta^{\dagger}$), and percentage of simulated instances with FDR $< 0.05$ and $\beta < 0.05$ ($\theta^{\ddagger}$) for each of the ten DE tests applied to data simulated from the Zhou dataset (30 instances) at three different sample sizes. Standard deviation in parentheses.

| Method | Sample size per group | | |
|---|---|---|---|
| | 30 | 50 | 100 |
| SIEVEseq | 0.014 (0.007) | 0.011 (0.009) | 0.012 (0.009)* |
| | 0.174 (0.034) | 0.056 (0.017) | 0.009 (0.006)† |
| | 0% | 36.7% | 100%‡ |
| ALDEx2 | 0.033 (0.013) | 0.034 (0.015) | 0.038 (0.014) |
| | 0.162 (0.030) | 0.053 (0.020) | 0.011 (0.007) |
| | 0% | 33.3% | 83.3% |
| NOISeq | 0.012 (0.012) | 0.008 (0.008) | 0.002 (0.004) |
| | 0.486 (0.051) | 0.257 (0.034) | 0.097 (0.024) |
| | 0% | 0% | 0% |
| edgeR | 0.028 (0.011) | 0.026 (0.014) | 0.030 (0.012) |
| | 0.082 (0.026) | 0.020 (0.010) | 0.005 (0.006) |
| | 10% | 93.3% | 93.3% |
| DESeq2 | 0.062 (0.015) | 0.049 (0.018) | 0.051 (0.015) |
| | 0.077 (0.024) | 0.023 (0.010) | 0.009 (0.007) |
| | 3.3% | 50% | 46.7% |
| voom | 0.044 (0.014) | 0.042 (0.016) | 0.044 (0.014) |
| | 0.106 (0.028) | 0.027 (0.012) | 0.005 (0.004) |
| | 0% | 70% | 63.3% |
| DSS | 0.052 (0.016) | 0.046 (0.017) | 0.052 (0.017) |
| | 0.068 (0.024) | 0.017 (0.008) | 0.005 (0.006) |
| | 10% | 60% | 43.3% |
| dearseq | 0.059 (0.015) | 0.049 (0.018) | 0.048 (0.017) |
| | 0.082 (0.028) | 0.021 (0.011) | 0.007 (0.006)† |
| | 0% | 53.3% | 60% |
| Wilcoxon | 0.047 (0.013) | 0.039 (0.016) | 0.042 (0.014) |
| | 0.096 (0.027) | 0.018 (0.009) | 0.002 (0.003) |
| | 3.3% | 73.3% | 66.7% |
| tweeDEseq | 0.049 (0.014) | 0.040 (0.015) | 0.044 (0.015) |
| | 0.096 (0.027) | 0.022 (0.010) | 0.008 (0.007) |
| | 3.3% | 66.7% | 66.7% |

**Table S3** Mean computing times (in seconds) for each of the ten DE tests applied to data simulated from the Valentim dataset[§], Kelmer dataset[¶], and Zhou dataset[£] (30 instances). Standard deviation in parentheses.

| Method | Sample size per group | | |
|---|---|---|---|
| | 30 | 50 | 100 |
| SIEVEseq | 125 (6) | 41 (3) | 37 (1)[§] |
| | 121 (6) | 38 (3) | 32 (1)[¶] |
| | 121 (6) | 41 (3) | 36 (1)[£] |
| ALDEx2 | 34 (0.4) | 26 (0.6) | 46 (0.7) |
| | 32 (0.3) | 26 (0.3) | 45 (0.7) |
| | 33 (0.4) | 26 (0.4) | 45 (0.4) |
| NOISeq | 5.4 (0.12) | 7.3 (0.14) | 11.6 (0.04) |
| | 5.4 (0.15) | 7.1 (0.14) | 11.5 (0.22) |
| | 5.4 (0.16) | 7.2 (0.20) | 11.6 (0.13) |
| edgeR | 0.6 (0.01) | 0.9 (0.02) | 1.6 (0.02) |
| | 0.6 (0.01) | 0.9 (0.01) | 1.6 (0.02) |
| | 0.6 (0.01) | 0.9 (0.01) | 1.6 (0.02) |
| DESeq2 | 2.2 (0.04) | 3.0 (0.06) | 5.0 (0.03) |
| | 2.1 (0.02) | 3.0 (0.07) | 5.0 (0.01) |
| | 2.2 (0.09) | 3.0 (0.02) | 5.1 (0.11) |
| voom | 0.7 (0.01) | 1.1 (0.02) | 1.9 (0.02) |
| | 0.7 (0.01) | 1.1 (0.08) | 1.9 (0.02) |
| | 0.7 (0.01) | 1.1 (0.01) | 1.9 (0.01) |
| DSS | 1.6 (0.01) | 2.4 (0.01) | 4.7 (0.04) |
| | 1.7 (0.01) | 2.6 (0.02) | 5.0 (0.04) |
| | 1.6 (0.02) | 2.5 (0.02) | 4.7 (0.03) |
| dearseq | 0.3 (0.02) | 0.5 (0.07) | 1.0 (0.02) |
| | 0.3 (0.01) | 0.5 (0.02) | 1.0 (0.02) |
| | 0.3 (0.02) | 0.5 (0.02) | 1.0 (0.02) |
| Wilcoxon | 1.4 (0.07) | 1.9 (0.07) | 2.8 (0.10) |
| | 1.4 (0.08) | 1.9 (0.02) | 2.7 (0.04) |
| | 1.4 (0.02) | 1.9 (0.03) | 2.7 (0.03) |
| tweeDEseq | 549 (21) | 566 (18) | 639 (17) |
| | 565 (26) | 590 (33) | 609 (27) |
| | 521 (19) | 529 (13) | 594 (16) |

**Table S4** The mean of false discovery rate (FDR*) and mean probability of Type II error ($\beta^\dagger$) for SIEVEseq applied to data simulated from the Valentim, Kelmer and Zhou datasets (30 instances) at three different sample sizes with different pseudo-values. Standard deviation in parentheses.

| Dataset | Pseudo-value | Sample size per group | | |
|---|---|---|---|---|
| | | 30 | 50 | 100 |
| Valentim | 0.01 | 0.031 (0.015) | 0.038 (0.012) | 0.042 (0.014)* |
| | | 0.228 (0.026) | 0.083 (0.019) | 0.027 (0.012)† |
| | 0.1 | 0.017 (0.012) | 0.016 (0.007) | 0.017 (0.009) |
| | | 0.226 (0.028) | 0.076 (0.019) | 0.016 (0.009) |
| | 0.5 | 0.014 (0.011) | 0.012 (0.006) | 0.012 (0.007) |
| | | 0.221 (0.028) | 0.066 (0.015) | 0.008 (0.007) |
| Kelmer | 0.01 | 0.026 (0.010) | 0.033 (0.012) | 0.033 (0.017) |
| | | 0.010 (0.007) | 0.007 (0.006) | 0.002 (0.003) |
| | 0.1 | 0.018 (0.008) | 0.018 (0.009) | 0.017 (0.012) |
| | | 0.008 (0.006) | 0.004 (0.005) | 0.002 (0.003) |
| | 0.5 | 0.016 (0.009) | 0.013 (0.007) | 0.013 (0.009) |
| | | 0.005 (0.004) | 0.003 (0.004) | 0.001 (0.002) |
| Zhou | 0.01 | 0.036 (0.015) | 0.051 (0.019) | 0.060 (0.019) |
| | | 0.170 (0.034) | 0.066 (0.019) | 0.020 (0.010) |
| | 0.1 | 0.021 (0.011) | 0.018 (0.014) | 0.018 (0.010) |
| | | 0.173 (0.034) | 0.060 (0.019) | 0.014 (0.008) |
| | 0.5 | 0.014 (0.007) | 0.011 (0.009) | 0.012 (0.009) |
| | | 0.172 (0.034) | 0.056 (0.017) | 0.009 (0.006) |

**Table S5** The mean false discovery rate (FDR*), mean probability of Type II error ($\beta^{\dagger}$), and percentage of simulated instances with FDR $< 0.05$ and $\beta < 0.2$ ($\theta^{\ddagger}$) for each of the nine DE tests applied to data simulated from the Dolgalev dataset (30 instances) at two different sample sizes. Standard deviations are shown in parentheses.

| Method | Sample size per group | |
|---|---|---|
| | 30 | 50 |
| SIEVEseq | 0.016 (0.020)<br>0.398 (0.117)<br>3.3% | 0.025 (0.045)*<br>0.150 (0.043)$^{\dagger}$<br>83.3%$^{\ddagger}$ |
| ALDEx2 | 0.005 (0.010)<br>0.493 (0.110)<br>0% | 0.022 (0.048)<br>0.204 (0.048)<br>50.0% |
| NOISeq | 0.009 (0.014)<br>0.749 (0.076)<br>0% | 0.023 (0.049)<br>0.599 (0.068)<br>0% |
| edgeR | 0.197 (0.074)<br>0.254 (0.104)<br>0% | 0.231 (0.131)<br>0.114 (0.046)<br>0% |
| DESeq2 | 0.153 (0.072)<br>0.312 (0.109)<br>0% | 0.179 (0.129)<br>0.170 (0.059)<br>0% |
| voom | 0.034 (0.027)<br>0.294 (0.099)<br>10.0% | 0.069 (0.110)<br>0.094 (0.034)<br>60.0% |
| DSS | 0.126 (0.072)<br>0.403 (0.132)<br>0% | 0.186 (0.128)<br>0.194 (0.067)<br>0% |
| dearseq | 0.034 (0.029)<br>0.529 (0.120)<br>0% | 0.063 (0.106)<br>0.319 (0.075)<br>0% |
| tweeDEseq | 0.030 (0.026)<br>0.540 (0.120)<br>0% | 0.062 (0.093)<br>0.303 (0.071)<br>3.3% |

**Table S6** The mean false discovery rate (FDR*), mean probability of Type II error ($\beta^{\dagger}$), and percentage of simulated instances with FDR $< 0.05$ and $\beta < 0.2$ ($\theta^{\ddagger}$) for each of the ten DE tests applied to data simulated from the Furusawa dataset (30 instances) at two different sample sizes. Standard deviations are shown in parentheses.

| Method | Sample size per group | |
|---|---|---|
| | 30 | 50 |
| SIEVEseq | 0.012 (0.014)<br>0.239 (0.077)<br>26.7% | 0.020 (0.028)*<br>0.106 (0.069)†<br>76.7%‡ |
| ALDEx2 | 0.007 (0.010)<br>0.250 (0.080)<br>30.0% | 0.027 (0.045)<br>0.108 (0.061)<br>70.0% |
| NOISeq | 0.016 (0.016)<br>0.587 (0.108)<br>0% | 0.017 (0.015)<br>0.391 (0.116)<br>0% |
| edgeR | 0.068 (0.035)<br>0.159 (0.087)<br>16.7% | 0.131 (0.083)<br>0.064 (0.039)<br>10.0% |
| DESeq2 | 0.083 (0.039)<br>0.147 (0.079)<br>16.7% | 0.138 (0.088)<br>0.064 (0.037)<br>10.0% |
| voom | 0.023 (0.020)<br>0.172 (0.065)<br>66.7% | 0.053 (0.063)<br>0.067 (0.051)<br>66.7% |
| DSS | 0.084 (0.042)<br>0.182 (0.091)<br>16.7% | 0.140 (0.091)<br>0.071 (0.042)<br>13.3% |
| dearseq | 0.034 (0.028)<br>0.308 (0.083)<br>10.0% | 0.058 (0.069)<br>0.177 (0.073)<br>36.7% |
| tweeDEseq | 0.032 (0.030)<br>0.334 (0.078)<br>3.3% | 0.058 (0.069)<br>0.186 (0.071)<br>36.7% |

**Table S7** The number and percentages of DV, pure DS, and DVS genes (gene set size in parentheses) detected using SIEVEseq that are collaterally detected using 7 DE methods. Abbreviations: DVS = non-DE, DV, and DS; Union = the union set of pure DV, pure DS, and DVS genes; DE Union = the union set of DE genes detected using edgeR, DESeq2, voom, ALDEx2, tweeDEseq, DSS, and Wilcoxon.

| Gene set (size) | pure DV (1843) | pure DS (919) | DVS (99) | Union (2861) |
|---|---|---|---|---|
| edgeR (3942) | 16.3% (301/1843) | 7.5% (69/919) | 17.2% (17/99) | 13.5% (387/2861) |
| DESeq2 (4155) | 17.7% (327/1843) | 8.2% (75/919) | 20.2% (20/99) | 14.8% (422/2861) |
| voom (3636) | 5.9% (109/1843) | 5.3% (49/919) | 1.0% (1/99) | 5.6% (159/2861) |
| ALDEx2 (3230) | 2.2% (41/1843) | 3.2% (29/919) | 0.0% (0/99) | 2.4% (70/2861) |
| tweeDEseq (3553) | 11.1% (205/1843) | 4.8% (44/919) | 6.1% (6/99) | 8.9% (255/2861) |
| DSS (4171) | 17.5% (322/1843) | 8.1% (74/919) | 19.2% (19/99) | 14.5% (415/2861) |
| Wilcoxon (3330) | 3.2% (59/1843) | 3.6% (33/919) | 1.0% (1/99) | 3.3% (93/2861) |
| DE Union (4770) | 20.1% (371/1843) | 11.5% (106/919) | 20.2% (20/99) | 17.4% (497/2861) |

**Table S8** Summary of statistically significant results from WebGestalt (affinity propagation clustering; top 20 GO terms in biological processes) with the union of DE, DV, and DS genes as query gene list.

| GO ID | Description | Size | Expect | Ratio | $p$-value | FDR |
|---|---|---|---|---|---|---|
| GO:0072599 | establishment of protein localization to endoplasmic reticulum | 111 | 25.87 | 2.67 | <2.2e-16 | <2.2e-16 |
| GO:0022610 | biological adhesion | 1377 | 320.91 | 1.39 | 8.88e-16 | 1.14e-12 |
| GO:0010647 | positive regulation of cell communication | 1733 | 403.87 | 1.33 | 5.11e-15 | 4.04e-12 |
| GO:0001568 | blood vessel development | 654 | 152.41 | 1.55 | 3.85e-14 | 2.19e-11 |
| GO:0030198 | extracellular matrix organization | 347 | 80.87 | 1.77 | 5.57e-14 | 2.98e-11 |

**Table S9** Summary of statistically significant results from WebGestalt (affinity propagation clustering; top 20 GO terms in biological processes) with high confidence DE genes as query gene list.

| GO ID | Description | Size | Expect | Ratio | $p$-value | FDR |
|---|---|---|---|---|---|---|
| GO:0001568 | blood vessel development | 714 | 69.71 | 2.03 | 1.20e-16 | 4.51e-13 |
| GO:0031589 | cell-substrate adhesion | 350 | 34.17 | 2.52 | 3.48e-16 | 7.87e-13 |
| GO:0042060 | wound healing | 422 | 41.20 | 2.26 | 3.01e-14 | 3.02e-11 |
| GO:0070848 | response to growth factor | 707 | 69.03 | 1.85 | 2.50e-12 | 1.74e-9 |

**Table S10** Summary of statistically significant results from WebGestalt (affinity propagation clustering; top 20 GO terms in biological processes) with nonDE, DV and/or DS genes (011, 001, and 011 classes) as query gene list.

| GO ID | Description | Size | Expect | Ratio | $p$-value | FDR |
|---|---|---|---|---|---|---|
| GO:0090150 | establishment of protein localization to membrane | 313 | 37.82 | 2.60 | <2.2e-16 | <2.2e-16 |
| GO:0006614 | SRP-dependent cotranslational protein targeting to membrane | 95 | 11.48 | 4.62 | <2.2e-16 | <2.2e-16 |
| GO:0034613 | cellular protein localization | 1815 | 219.33 | 1.43 | 5.47e-12 | 3.11e-09 |
| GO:0006412 | translation | 613 | 74.08 | 1.80 | 5.88e-12 | 3.15e-09 |

**Table S11** Summary of statistically significant results from WebGestalt (affinity propagation clustering; top 20 GO terms in biological processes) with pure DV genes (010 class) as query gene list.

| GO ID | Description | Size | Expect | Ratio | $p$-value | FDR |
|---|---|---|---|---|---|---|
| GO:0090150 | establishment of protein localization to membrane | 313 | 24.49 | 3.06 | <2.2e-16 | <2.2e-16 |
| GO:0045047 | protein targeting to ER | 107 | 8.37 | 6.21 | <2.2e-16 | <2.2e-16 |
| GO:0006401 | RNA catabolic process | 341 | 26.68 | 2.81 | 1.11e-16 | 7.76e-14 |
| GO:0006412 | translation | 613 | 47.96 | 2.08 | 9.66e-13 | 5.17e-10 |

**Table S12** Summary of statistically significant results from WebGestalt (affinity propagation clustering; top 20 GO terms in biological processes) with pure DS genes (001 class) as query gene list.

| GO ID | Description | Size | Expect | Ratio | $p$-value | FDR |
|---|---|---|---|---|---|---|
| GO:0019884 | antigen processing and presentation of exogenous antigen | 129 | 4.95 | 2.83 | 0.0004 | 0.61 |
| GO:0019363 | pyridine nucleotide biosynthetic process | 108 | 4.14 | 2.90 | 0.0009 | 0.61 |
| GO:0019752 | carboxylic acid metabolic process | 960 | 36.81 | 1.52 | 0.0011 | 0.61 |
| GO:0045619 | regulation of lymphocyte differentiation | 162 | 6.21 | 2.41 | 0.0014 | 0.61 |
| GO:1901844 | regulation of cell communication by electrical coupling involved in cardiac conduction | 7 | 0.27 | 11.18 | 0.0017 | 0.61 |

### S4 Supplementary Figures

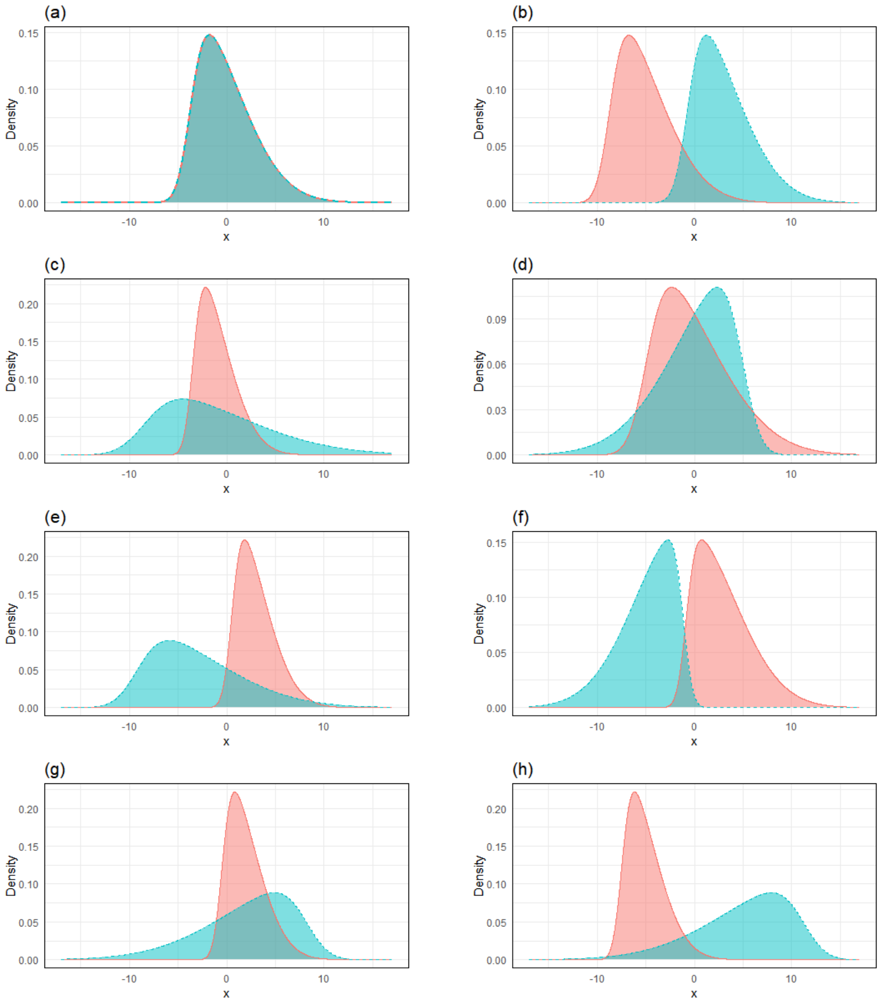


**Fig. S1** Examples of distribution of CLR-transformed data for two biological conditions (black and red) leading to the eight possible gene classes as determined by simultaneous testing of DE, DV, and DS. (a): non-DE, non-DV and non DS; (b): DE, non-DV and non-DS; (c): non-DE, DV and non-DS; (d): non-DE, non DV and DS; (e): DE, DV and non-DS; (f): DE, non-DV and DS; (g): non-DE, DV and DS; (h): DE, DV and DS.

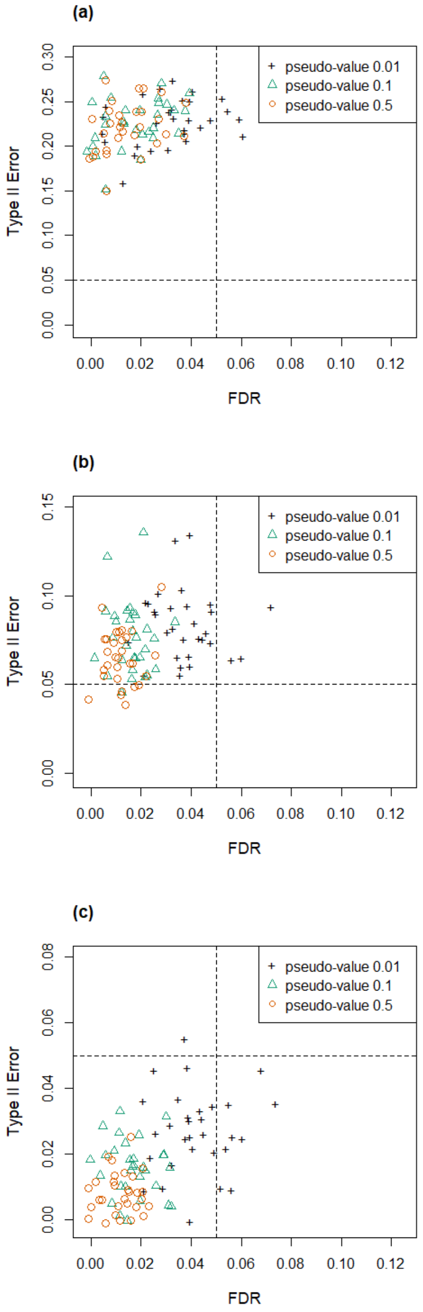


**Fig. S2** Scatter plots of the probability of Type II error ($\beta$) versus FDR for simulated data based on the Valentim dataset (30 instances), using the SIEVEseq DE test with pseudo-values of 0.01, 0.1, and 0.5, under three sample size scenarios per group: (a) 30; (b) 50; (c) 100. Dashed lines indicate the desired thresholds for FDR and $\beta$ of 0.05.

(a)

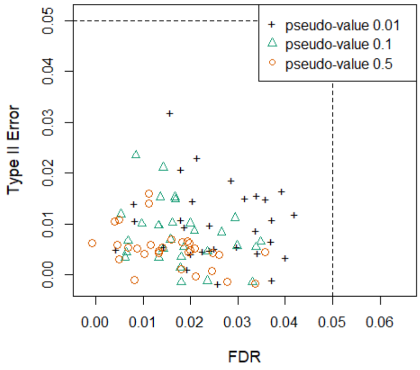


(b)

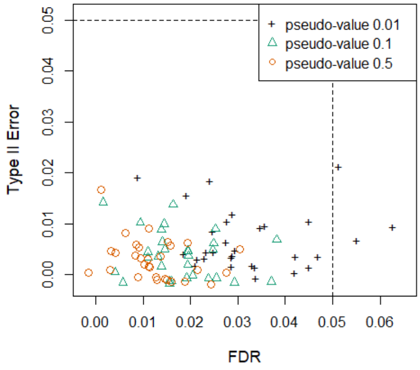


(c)

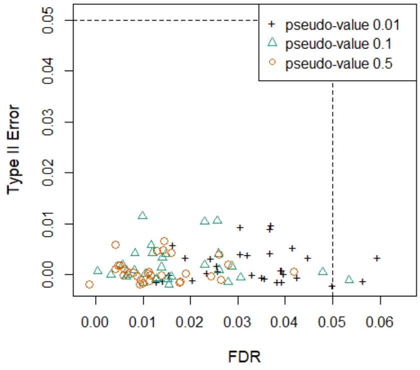


**Fig. S3** Scatter plots of the probability of Type II error ($\beta$) versus FDR for simulated data based on the Kelmer dataset (30 instances), using the SIEVEseq DE test with pseudo-values of 0.01, 0.1, and 0.5, under three sample size scenarios per group: (a) 30; (b) 50; (c) 100. Dashed lines indicate the desired thresholds for FDR and $\beta$ of 0.05.

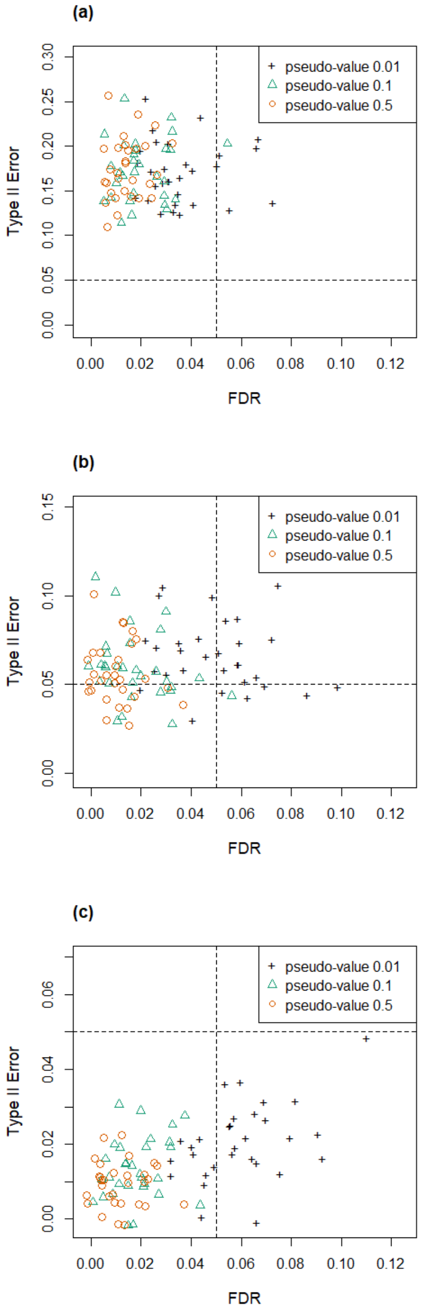


**Fig. S4** Scatter plots of the probability of Type II error ($\beta$) versus FDR for simulated data based on the Zhou dataset (30 instances), using the SIEVEseq DE test with pseudo-values of 0.01, 0.1, and 0.5, under three sample size scenarios per group: (a) 30; (b) 50; (c) 100. Dashed lines indicate the desired thresholds for FDR and $\beta$ of 0.05.

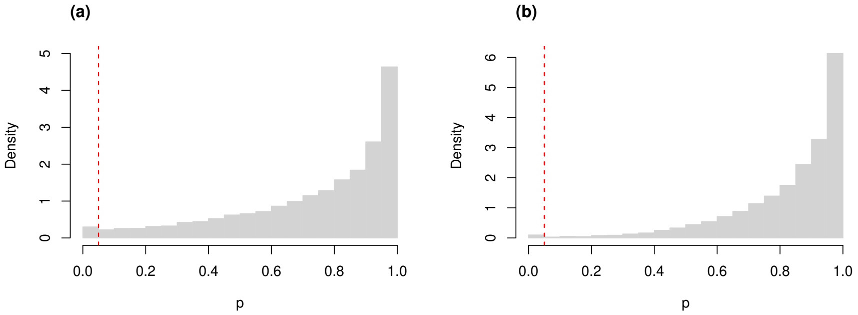


**Fig. S5** Histograms of the distribution of the $p$-values from Kolmogorov-Smirnov goodness-of-fit tests of the skew-normal model on genes from the (a) the control group ($p$-value $> 0.05$ for 98.5% of genes); and (b) the AD group ($p$-value $> 0.05$ for 99.5% of the genes). The red dashed lines mark the threshold $p$-value of 0.05.

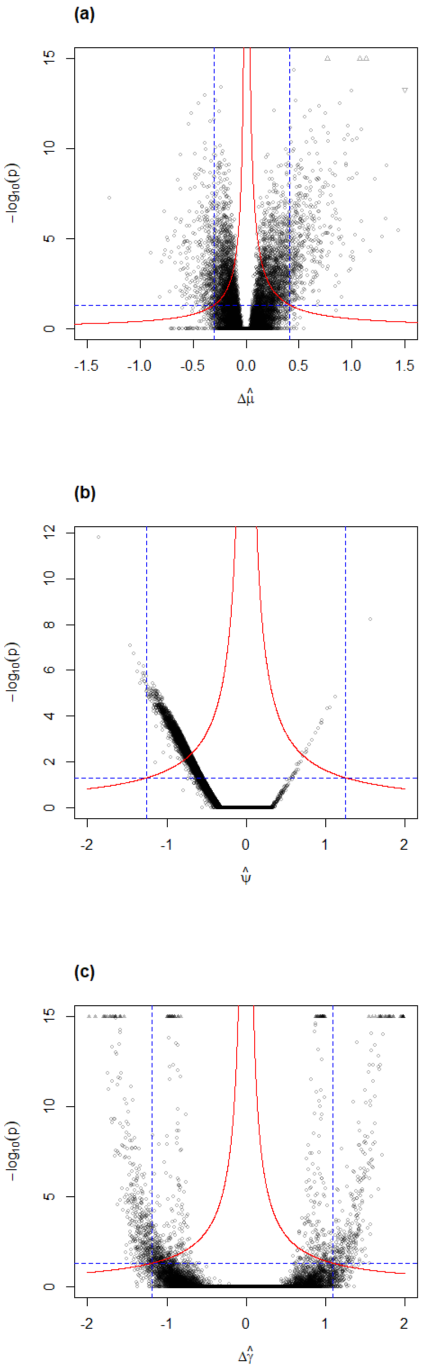


**Fig. S6** Volcano plots for DE, DV, and DS tests using SIEVEseq on the Mayo RNA-Seq dataset. (a) DE test (2773 DE genes); (b) DV test (2276 DV genes); and (c) DS test (1204 DS genes). Red lines indicate decision boundaries; blue lines indicate the threshold biological and statistical significance values used.

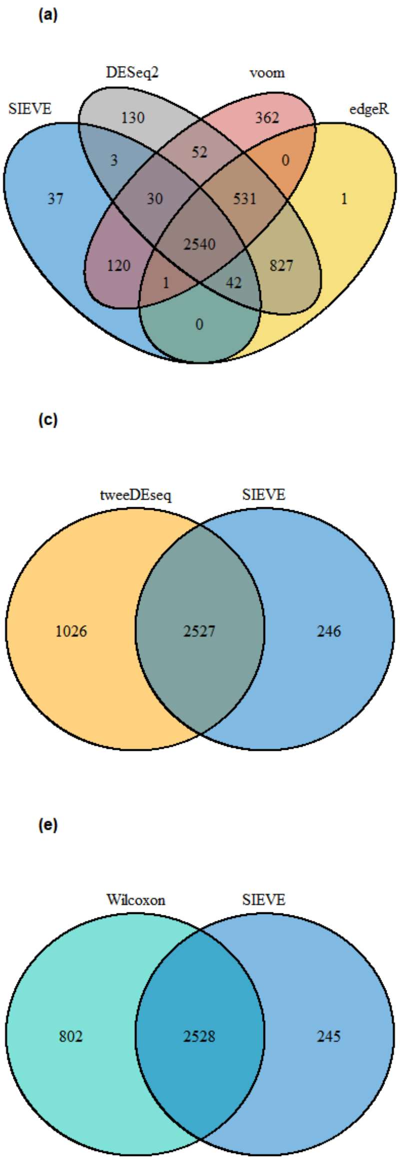


**Fig. S7** Venn diagrams of DE genes detected using SIEVEseq and (a) edgeR, DESeq2 and voom; (b) ALDEx2; (c) tweeDEseq; (d) DSS; (e) Wilcoxon; (f) NOISeq for the control vs. AD comparison.

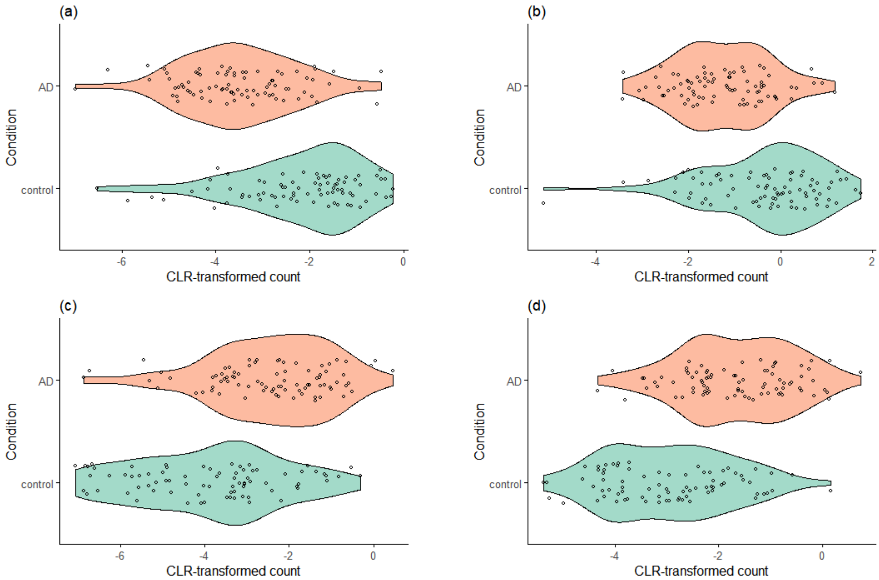


**Fig. S8** Violin plots of selected pure DE genes detected in the control vs. AD comparison. (a) *CRH*; (b) *SST*; (c) *S100A12*; and (d) *SERPINA5*.

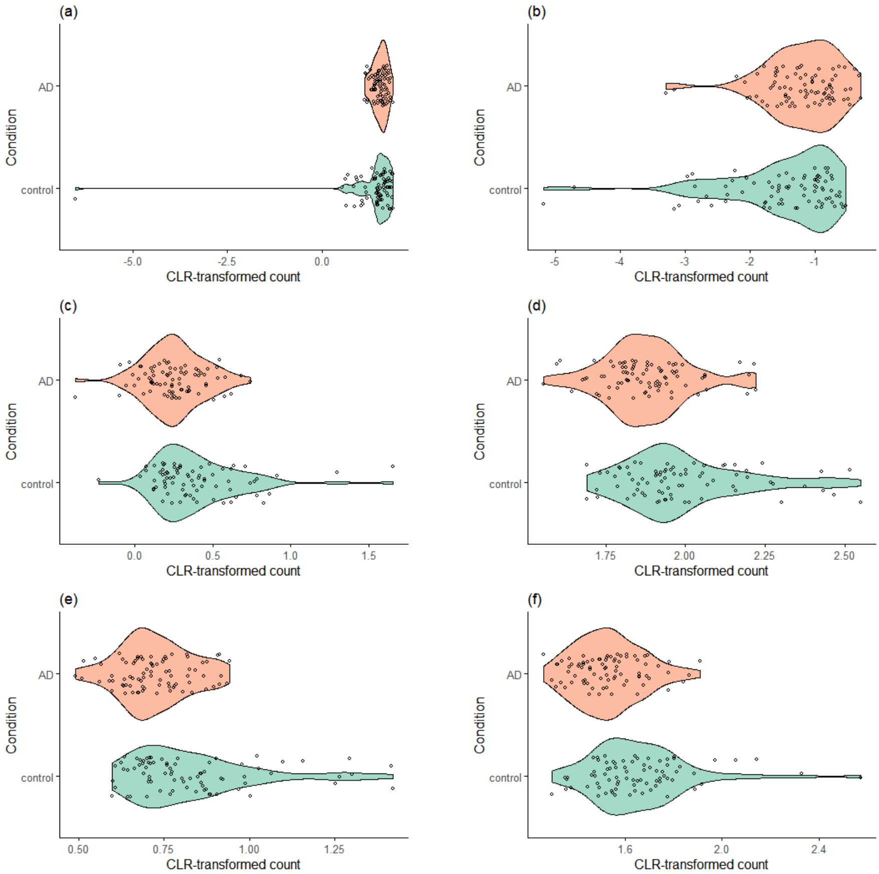


**Fig. S9** Violin plots of selected DE genes uniquely detected by SIEVEseq in the control vs. AD comparison. (a)*FOXG1*; (b) *HTR1E*; (c) *ZNF337*; (d) *EHMT2*; (e) *HARS2*; and (f) *TSC2*.

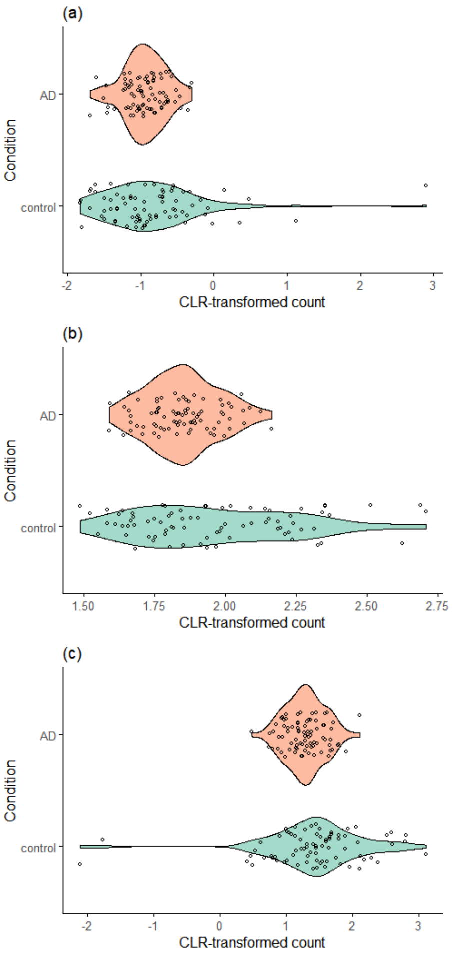


**Fig. S10** Violin plots of selected pure DV genes detected in the control vs. AD comparison. (a) *IL16*; (b) *SGTA*; and (c) *GP1BB*.

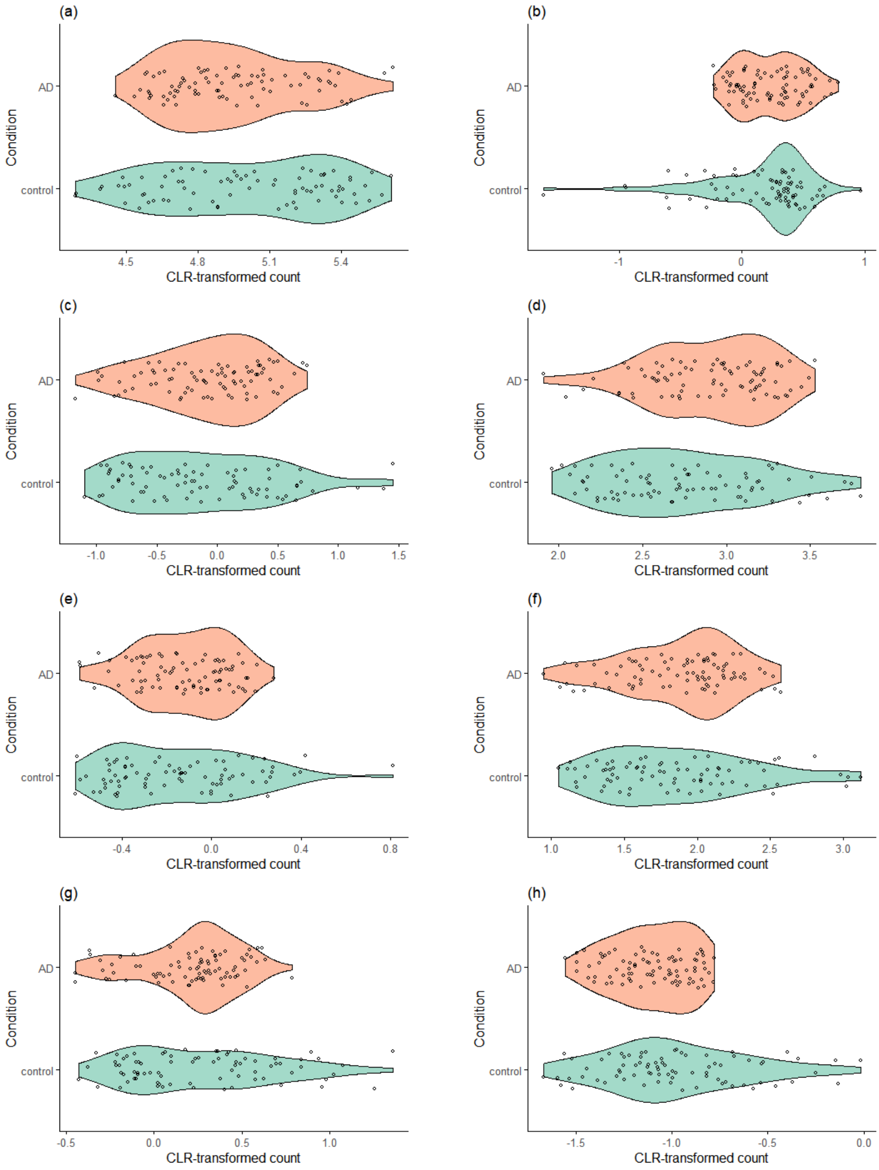


**Fig. S11** Violin plots of selected pure DS genes detected in the control vs. AD comparison. (a) *CALM1*; (b) *TSHZ3*; (c) *CYBA*; (d) *PLXNB1*; (e) *CLN3*; (f) *ADARB2*; (g) *MAN2B1*; and (h) *BACE1-AS*.

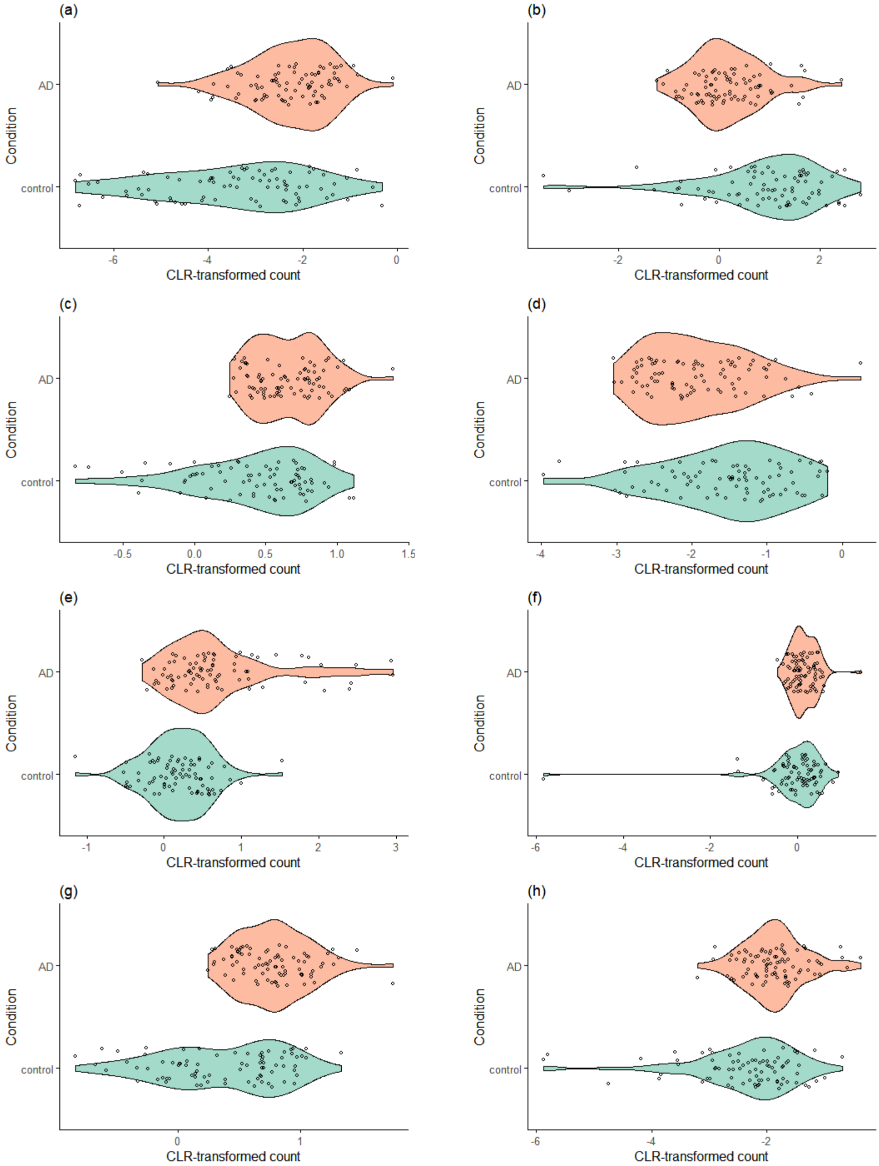


**Fig. S12** Violin plots of selected DEV, DES, and DEVS genes detected in the control vs. AD comparison. (a) *FCGR3B* (DEV gene), (b) *VGF* (DES gene), (c) *CPNE8* (DES gene), (d) *IGF1* (DES gene), (e) *LEPR* (DES gene), (f) *FEZF2* (DEVS gene), (g) *SOX5* (DEVS gene), and (h) *POSTN* (DEVS gene).

### S5 Biological Significance of Detected DE, DV, and DS Genes

*FMOD* encodes the fibromodulin protein which activates the complement pathway through direct binding to the C1q protein, a component of the C1 complex known to interact with abnormal protein structures in AD and prion diseases (Sjöberg et al, 2005).

Deficiency in the expression of *TTC7A* has been reported to produce mis-activation of the RhoA signaling pathway, leading to abberant proliferation, adhesion, and migratory capacities of lymphocytes (Lemoine et al, 2014). Interestingly, the RhoA/ROCK signaling pathway activation is known to regulate the production of amyloid beta (A$\beta$), formation of neurofibrillary tangles, and neuroinflammatory responses (Cai et al, 2021). In connection, the histamine receptor H1 (HRH1) protein responds to the histamine ligand by activating G$\alpha$q-mediated signaling pathway (van Unen et al, 2016), which activates RhoA (Chikumi et al, 2002).

The *CRH* gene encodes a member of the corticotropin-releasing factor family, which initiates neuroendocrine responses to stress by activating the hypothalamic-pituitary-adrenal (HPA) axis (Smith and Vale, 2006). In the 1980s, CRH-like immunoreactivity was found to be significantly decreased in the temporal, frontal and occipital cortex of AD patients, with reciprocal increase in CRH receptors in the same brain regions (De Souza et al, 1987).

The possible association of *MTMR7* with AD was only very recently known through a genome-wide association study, whereby the chromosome 8 locus on which *MTMR7* is located is shown to be associated with cognitive resilience among amyloid-Positron Emission Tomography (PET) positive subjects (Ramanan et al, 2021).

### S6 List of R Packages Used

The following R packages (in alphabetical order) were used in the present work: `ALDEx2` (Fernandes et al, 2014), `compositions` (van den Boogaart et al, 2022), `dearseq` (Gauthier et al, 2020), `DESeq2` (Love et al, 2014), `dplyr` (Wickham et al, 2022a), `DSS` (Wu et al, 2013), `edgeR` (Robinson et al, 2010), `fgsea` (Korotkevich et al, 2019), `ggplot2` (Wickham, 2016), `gplots` (Warnes et al, 2022), `gridExtra` (Baptiste, 2017), `httr` (Wickham, 2022), `jsonlite` (Ooms, 2014), `limma` (Law et al, 2014), `MASS` (Venables and Ripley, 2002), `NOISeq` (Tarazona et al, 2015), `org.Hs.eg.db` (Carlson, 2025), `polyester` (Frazee et al, 2022), `readr` (Wickham et al, 2022b), `SimSeq` (Benidt and Nettleton, 2015), `sn` (Azzalini, 2022), `tweeDEseq` (Esnaola et al, 2013), `VennDiagram` (Chen, 2022).

### S7 Captions for Online Supplementary Tables

Table S13: List of DE genes detected by SIEVEseq, edgeR, DESeq2, limma-voom, ALDEx2, tweeDEseq, DSS, Wilcoxon rank-sum test, and NOISeq for the control vs. AD comparison in the analysis of the Mayo RNA-Seq dataset. Available at https://github.com/Divo-Lee/SIEVEseq.

Table S14: List of DE, DV, and DS genes identified by SIEVEseq for the control vs. AD comparison in the analysis of the Mayo RNA-Seq dataset. Available at https://github.com/Divo-Lee/SIEVEseq.

### S8 Acknowledgements

The materials presented here are based on the contents of the PhD thesis of HL.

### References

Azzalini A (1985) A class of distributions which includes the normal ones. Scandinavian Journal of Statistics 12(2):171–178

Azzalini A (2022) The R package `sn`: The skew-normal and related distributions such as the skew-$t$ and the SUN (version 2.1.0). Università degli Studi di Padova, Italia, URL https://cran.r-project.org/package=sn, home page: http://azzalini.stat.unipd.it/SN/

Azzalini A, Arellano-Valle RB (2013) Maximum penalized likelihood estimation for skew-normal and skew-t distributions. Journal of Statistical Planning and Inference 143(2):419–433

Azzalini A, Capitanio A (2014) The Skew-Normal and Related Families. Cambridge University Press, Cambridge

Baptiste A (2017) gridExtra: Miscellaneous Functions for "Grid" Graphics. URL https://CRAN.R-project.org/package=gridExtra, R package version 2.3

Benidt S, Nettleton D (2015) Simseq: a nonparametric approach to simulation of RNA-sequence datasets. Bioinformatics 31(13):2131–2140

Cabras S, Racugno W, Castellanos ME, et al (2012) A matching prior for the shape parameter of the skew-normal distribution. Scandinavian Journal of Statistics 39(2):236–247

Cai R, Wang Y, Huang Z, et al (2021) Role of RhoA/ROCK signaling in Alzheimer's disease. Behavioural Brain Research 414:113,481

Carlson M (2025) org.Hs.eg.db: Genome wide annotation for Human. R package version 3.21.0

Chen H (2022) VennDiagram: Generate High-Resolution Venn and Euler Plots. URL https://CRAN.R-project.org/package=VennDiagram, R package version 1.7.3

Chikumi H, Vázquez-Prado J, Servitja J, et al (2002) Potent activation of Rhoa by G$\alpha$q and Gq-coupled receptors. Journal of Biological Chemistry

277(30):27,130–27,134

De Souza EB, Whitehouse PJ, Price DL, et al (1987) Abnormalities in corticotropin-releasing hormone (CRH) in Alzheimer's disease and other human disorders. Annals of the New York Academy of Sciences 512(1):237–247

Esnaola M, Puig P, Gonzalez D, et al (2013) A flexible count data model to fit the wide diversity of expression profiles arising from extensively replicated RNA-seq experiments. BMC Bioinformatics 14:254

Fernandes AD, Reid JN, Macklaim JM, et al (2014) Unifying the analysis of high-throughput sequencing datasets: characterizing RNA-seq, 16S rRNA gene sequencing and selective growth experiments by compositional data analysis. Microbiome 2:15

Frazee AC, Jaffe AE, Kirchner R, et al (2022) polyester: Simulate RNA-seq reads. R package version 1.32.0

Gauthier M, Agniel D, Thiébaut R, et al (2020) dearseq: a variance component score test for RNA-seq differential analysis that effectively controls the false discovery rate. NAR Genomics and Bioinformatics 2(4):lqaa093

Korotkevich G, Sukhov V, Budin N, et al (2019) Fast gene set enrichment analysis. bioRxiv https://doi.org/10.1101/060012, URL http://biorxiv.org/content/early/2016/06/20/060012

Law C, Chen Y, Shi W, et al (2014) voom: precision weights unlock linear model analysis tools for RNA-seq read counts. Genome Biology 15:R29

Lemoine R, Pachlopnik-Schmid J, Farin H, et al (2014) Immune deficiency-related enteropathy-lymphocytopenia-alopecia syndrome results from tetratricopeptide repeat domain 7A deficiency. Journal of Allergy and Clinical Immunology 134(6):1354–1364

Love M, Huber W, Anders S (2014) Moderated estimation of fold change and dispersion for RNA-seq data with DESeq2. Genome Biology 15:550

O'Hagan A, Leonard T (1976) Bayes estimation subject to uncertainty about parameter constraints. Biometrika 63(1):201–203

Ooms J (2014) The jsonlite package: A practical and Consistent mapping between JSON data and R objects. arXiv:14032805 [statCO] URL https://arxiv.org/abs/1403.2805

Ramanan VK, Lesnick TG, Przybelski SA, et al (2021) Coping with brain amyloid: genetic heterogeneity and cognitive resilience to Alzheimer's pathophysiology. Acta Neuropathologica Communications 9:48

Robinson MD, McCarthy DJ, Smyth GK (2010) edgeR: a Bioconductor package for differential expression analysis of digital gene expression data. Bioinformatics 26(1):139–140

Sjöberg A, Önnerfjord P, Mörgelin M, et al (2005) The extracellular matrix and inflammation: fibromodulin activates the classical pathway of complement by directly binding C1q. Journal of Biological Chemistry 280(37):32,301–32,308

Smith SM, Vale WW (2006) The role of the hypothalamic-pituitary-adrenal axis in neuroendocrine responses to stress. Dialogues in Clinical Neuroscience 8(4):383–395

Tarazona S, Furió-Tarí P, Turrà D, et al (2015) Data quality aware analysis of differential expression in RNA-seq with NOISeq R/Bioc package. Nucleic Acids Research 43(21):e140

van Unen J, Rashidfarrokhi A, Hoogendoorn E, et al (2016) Quantitative single-cell analysis of signaling pathways activated immediately downstream of histamine receptor subtypes. Molecular Pharmacology 90(3):162–176

van den Boogaart KG, Tolosana-Delgado R, Bren M (2022) compositions: Compositional Data Analysis. URL https://CRAN.R-project.org/package=compositions, R package version 2.0-4

Venables WN, Ripley BD (2002) Modern Applied Statistics with S, 4th edn. Springer, New York, URL https://www.stats.ox.ac.uk/pub/MASS4/

Warnes GR, Bolker B, Bonebakker L, et al (2022) gplots: Various R Programming Tools for Plotting Data. URL https://CRAN.R-project.org/package=gplots, R package version 3.1.3

Wickham H (2016) ggplot2: Elegant Graphics for Data Analysis. Springer-Verlag New York, URL https://ggplot2.tidyverse.org

Wickham H (2022) httr: Tools for Working with URLs and HTTP. URL https://CRAN.R-project.org/package=httr, R package version 1.4.4

Wickham H, François R, Henry L, et al (2022a) dplyr: A Grammar of Data Manipulation. URL https://CRAN.R-project.org/package=dplyr, R package version 1.0.10

Wickham H, Hester J, Bryan J (2022b) readr: Read Rectangular Text Data. URL https://CRAN.R-project.org/package=readr, R package version 2.1.2

Wu H, Wang C, Wu Z (2013) A new shrinkage estimator for dispersion improves differential expression detection in RNA-seq data. Biostatistics 14(2):232–243

Xiao Y, Hsiao TH, Suresh U, et al (2014) A novel significance score for gene selection and ranking. Bioinformatics 30(6):801–807
